# HIV integration in the human brain is linked to microglial activation and 3D genome remodeling

**DOI:** 10.1101/2022.05.03.490485

**Authors:** Amara L. Plaza-Jennings, Aditi Valada, Callan O’Shea, Marina Iskhakova, Benxia Hu, Behnam Javidfar, Gabriella Ben Hutta, Tova Lambert, Jacinta Murray, Bibi Kassim, Sandhya Chandrasekaran, Benjamin K. Chen, Susan Morgello, Hyejung Won, Schahram Akbarian

## Abstract

Exploration of genome organization and function in the HIV infected brain is critical to aid in the development of treatments for HIV-associated neurocognitive disorder (HAND) and HIV cure strategies. Here, we generated a resource comprised of single nuclei transcriptomics, complemented by cell-type-specific Hi-C chromosomal conformation (‘3D genome’) and viral integration site sequencing (IS-seq) in frontal brain tissues from individuals with HIV encephalitis (HIVE), HIV-infected people without encephalitis (HIV+), and HIV uninfected (HIV-) controls. We observed profound 3D genomic reorganization of open/repressive (A/B) compartment structures encompassing 6.4% of the HIVE microglial genome that was associated with transcriptomic reprogramming, including down-regulation of homeostasis and synapse-related functions and robust activation of interferon signaling and cell migratory pathways. HIV RNA was detected in 0.003% of all nuclei in HIVE brain, predominantly in the most activated microglia where it ranked as the second most highly expressed transcript. Microglia from HIV+ brains showed, to a lesser extent, similar transcriptional alterations. IS-seq recovered 1,221 insertion events in glial nuclei that were enriched for chromosomal domains newly mobilized into a permissive chromatin environment in HIVE microglia. Brain and peripheral myeloid cell integration revealed a preference overall for transcription-permissive chromatin, but robust differences in the frequency of recurrent insertions, intergenic integration, and enrichment for pre-integration complex-associated factors at integration sites. Our resource highlights critical differences in the genomic patterns of HIV infection in brain versus blood and points to a dynamic interrelationship between inflammation-associated 3D genome remodeling and successful integration in brain.

## INTRODUCTION

HIV enters the brain within the first two weeks of infection, and biomarkers of central nervous system (CNS) inflammation as well as neurologic symptoms have been observed in acute HIV disease (*1–3*). Considering that roughly 7x10^9^ microglial cells, the primary CNS cell type infected by HIV, reside in an adult human brain (*4–7*), the central nervous system is potentially a large reservoir site (*2, 8*). Additionally, the brain is one of the organs with the highest burden of HIV-associated disease. HIV-associated neurocognitive disorder (HAND) affects 20-50% of the 37 million people living with HIV (PLWH), with the milder forms of HAND predominating in the era of combined antiretroviral therapy (cART) (*9–11*).

Therefore, genomic and transcriptomic exploration of the HIV-infected human brain is critical for both the understanding and development of treatments for HAND and HIV cure strategies. However, to date, genomic studies in the HIV infected brain have primarily focused on bulk tissue gene expression profiling and demonstrated neuroinflammation and metabolic alterations (*12, 13*). Herein we describe the first cell-type-specific, integrative genomics studies of frontal lobe tissues from individuals who were HIV-infected with encephalitis (HIVE), HIV-infected without encephalitis (HIV+), and HIV-uninfected (HIV-). We focused on HIVE as a model of active brain infection and profiled HIV integration patterns, chromosomal conformation (‘3D genome’) mapping, and single nucleus transcription. We uncovered large-scale rearrangements of the HIVE microglial 3D genome affecting hundreds of megabases, linked to microglial transcriptome reprogramming defined by strong immune activation signatures and loss of homeostatic support functions for the neuronal synapse. We observed preferential viral integration in regions of HIVE 3D genomic restructuring and preferential viral transcription in activated microglia, suggesting that microglial activation potentiates infection. Our integrative study, generating some of the first microglia-specific 3D genome maps and single cell transcriptome profiles in the context of neuroinflammation, provides an important and valuable new resource to the field. Our discovery that chromosomal loci defined by 3D mobilization into a more permissive chromatin environment show increased risk for viral integration inside the microglial nucleus strongly points to a critical role of inflammation-driven reorganization of the chromosomal conformations for persistent HIV infection.

## RESULTS

### Subhead 1: Transcriptomic profiling of the HIV-infected brain at a single nucleus resolution

Using dual-labeling RNA fluorescence in situ hybridization (RNA FISH) for HIV transcripts and immunohistochemistry for microglial markers, we first confirmed high levels of HIV RNA expression in encephalitic brain that showed a high degree of overlap with Iba1 labeled microglia (Fig. 1A). Next, in order to obtain deeper, cell-type specific resolution of HIV-infected and uninfected brain, we performed 10X Chromium single nucleus RNA-sequencing (snRNA-seq) of frontal lobe, including cortical gray matter (FC) and underlying subcortical white matter (WM), of HIV uninfected (HIV-, n=3), HIV infected (HIV+, n=3) and HIV infected with encephalitis (HIVE, n=7) samples (Fig. 1B, Table S1). We profiled a total of 69,843 nuclei to an average depth of 119,797 ± 32,766 reads and a median of 2,401 ± 221 expressed genes/nucleus (Data file S1). Consistent with previous studies (*14, 15*), we identified 17 cell type clusters, including 7 excitatory (Exc1-7) and 3 inhibitory neuronal subtypes (Inh1-3), oligodendrocytes (Ol), oligodendrocyte progenitor cells (OPC), astrocytes (Ast), endothelial cells (Endo), lymphoid cells (Lymph), microglia/macrophages (Mg), and immune oligodendroglia (ImOl). ImOl were a rare cell population of only 336 nuclei total with marker gene expression of both oligodendrocyte and immune-related genes that closely matched ImOl cells previously described in snRNA-seq profiling from subcortical white matter (Fig. 1C-D, for marker gene expression see Data file S2) (*16, 17*). We observed expression of CD163 in the Mg cluster, indicating that this cluster contained perivascular macrophages as well as microglia (Fig. 1D).

**Figure 1.**
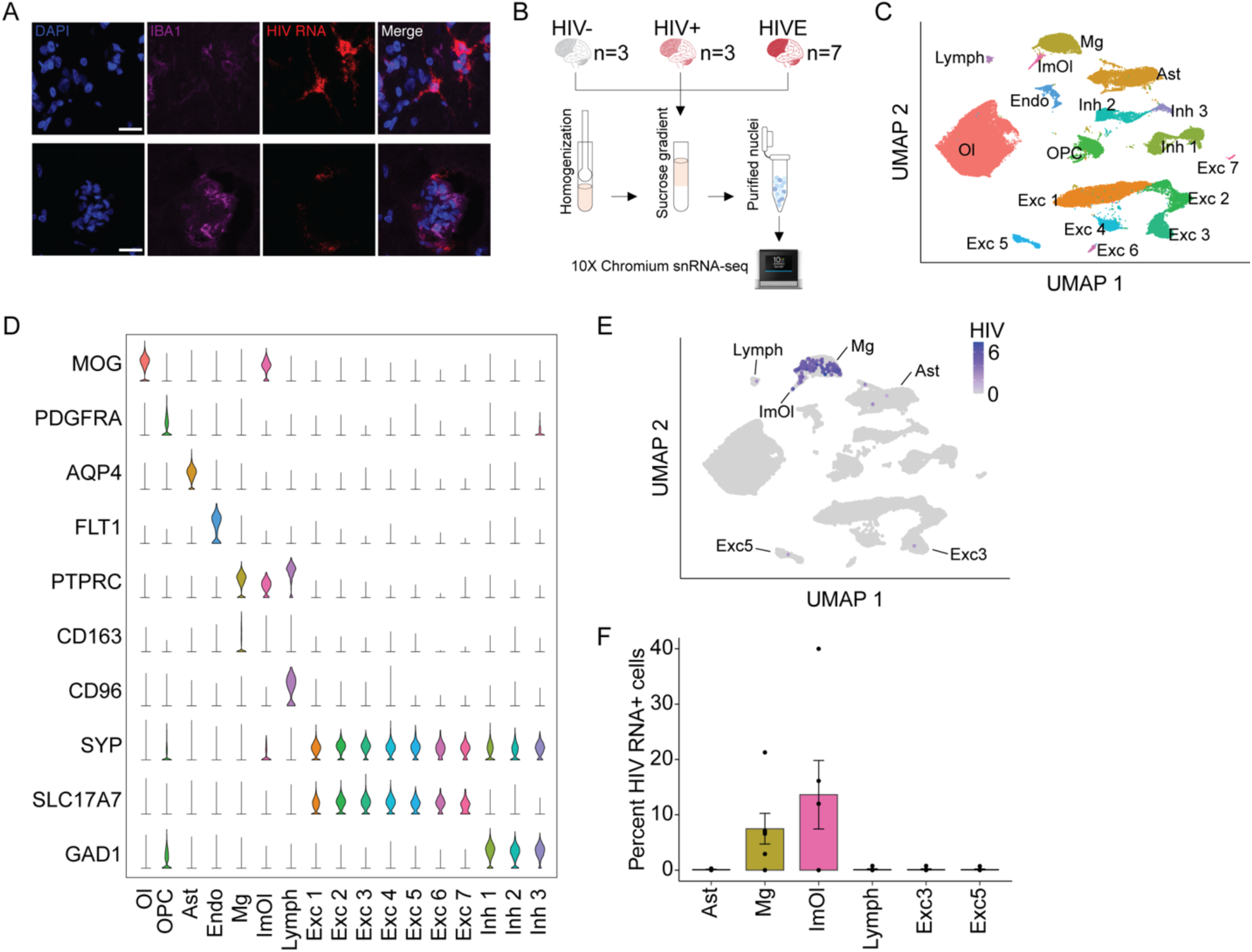
Cell type specificity and prevalence of HIV RNA+ nuclei in HIVE. (**A**) HIV RNA FISH in a histological section from a HIVE with immunofluorescence labeling of Iba1 microglia and DAPI nuclear stain. (**B**) Sample number and schematic preparation of tissue for 10X Chromium single nucleus RNA-sequencing. (**C**) Uniform Manifold Approximation and Projection (UMAP) plot showing clustering of identified cell clusters for n=69,843 total nuclei. Excitatory neurons subtypes 1-7 (Exc1-7), inhibitory neurons subtypes 1-3 (Inh1-3), oligodendrocytes (Ol), oligodendrocyte progenitor cells (OPC), endothelial cells (Endo), astrocytes (Ast), lymphoid cells (Lymph), microglia (Mg), and immune oligodendroglia (ImOl). (**D**) Marker gene expression for each cluster. Each row is scaled to the maximum of expression for that gene. (**E**) UMAP plot displaying log normalized expression of HIV transcripts. F) Percent of cells that express HIV RNA in each cluster (mean ± S.E.M) shown only for clusters that had any HIV RNA+ cells. Points represent individual donors.

We identified high confidence HIV RNA^+^ nuclei, using a stringent read count threshold determined by intermixing of HIVE and naïve mouse brain tissue controls for snRNA-seq (Fig. S1, see also Methods). We detected viral transcription in the Mg, ImOl, Lymph, Ast, Exc3, and Exc5 clusters, with the majority of HIV RNA+ cells in the Mg cluster (Fig. 1E-F, Data file S3). Overall, 4.8% of nuclei in the HIVE Mg cluster (133/2,756) and in HIVE ImOl (10/209) were HIV RNA^+^, while in comparison, the proportion of astrocytes, lymphocytes, and neurons harboring HIV RNA-expressing nuclei was lower by at least one order of magnitude (∼0.1 – 0.3%) with only 1-3 HIV RNA+ nuclei (Fig. 1E-F, Data file S3). Consistent with these findings, *CD4*, encoding the virus’ primary receptor (*18*), was only detectable in Mg and ImOl clusters, and furthermore, expression of *CCR5* and *CXCR4* transcripts, the two major viral co-receptors (*19*), was low overall across our different FC/WM cell types with the exception of robust *CXCR4* expression in the Lymph cluster (Fig. S2). Note that 87% (n=319) of all Lymph nuclei came from HIVE donors who had very low ante-mortem CD4 counts (Table S1). Additionally, HIV infections causes downregulation of *CD4* expression (*20*), thus very low *CD4* expression was observed in the Lymph cluster (Fig. S2). Finally, in contrast to more sensitive methods such as qPCR which was positive for HIV DNA in 3/3 HIV+ brains (Table S1), HIV transcription by single nuclei profiling was not detected in any nucleus from our N=3 HIV^+^ brains (Fig. S1B), while it was readily detectable in 5/7 HIVE brains (Data file S3).

### Subhead 2: Transcriptomic alterations in HIVE microglia are linked to large scale reorganization of chromosomal conformations

Cell culture models of acute immune activation have shown that peripheral immune cells undergo genomic responses to activating stimuli that include broad changes in chromosomal conformation together with dynamic alterations to gene expression activity, thereby profoundly affecting cellular function (*21–25*). However, despite these interesting findings from cultured lymphocytes and macrophages, genome reorganization upon an external stimulus has not been studied in microglia, and furthermore, 3D reorganization of any immune cell *in vivo* is completely unexplored.

Therefore, after having confirmed that microglia were the predominant cell type expressing HIV in the brain, we asked whether this cell type is, in the HIVE brain, characterized by distinct alterations in spatial genome organization. We generated genome-scale chromosomal contact maps for microglia and neurons from HIVE and HIV-brain (N=2 for each condition). For each brain, nuclei from FC/WM were extracted, fixed, and microglial and neuronal nuclei were isolated using fluorescence-activated nuclei sorting (FANS) with antibodies to the microglial-specific transcription factor Interferon regulatory factor 8 (Irf8) and the neuronal specific marker, neuronal nuclear protein (NeuN, Fig. 2A). We profiled 3D genome structure of Irf8+ and NeuN+ nuclei using *in situ* Hi-C, with samples sequenced to an average of >200M valid cis (intrachromosomal) chimeric reads (Data file S1). Cell-type specificity of sorted nuclei was confirmed by expression profiling of microglial and neuronal marker genes in RNA-seq from sorted nuclei (Fig. S3A). Next, we applied pairwise correlations to the cell type-specific Hi-C datasets and found that the largest difference was by cell type rather than diagnosis (Fig. S3B). However, among the microglial Hi-C libraries, we did observe stronger correlation within each diagnosis group than between cases and controls (Fig. S3B), strongly suggesting viral infection further modifies the Hi-C chromosomal contact map on a genome-wide scale.

**Figure 2.**
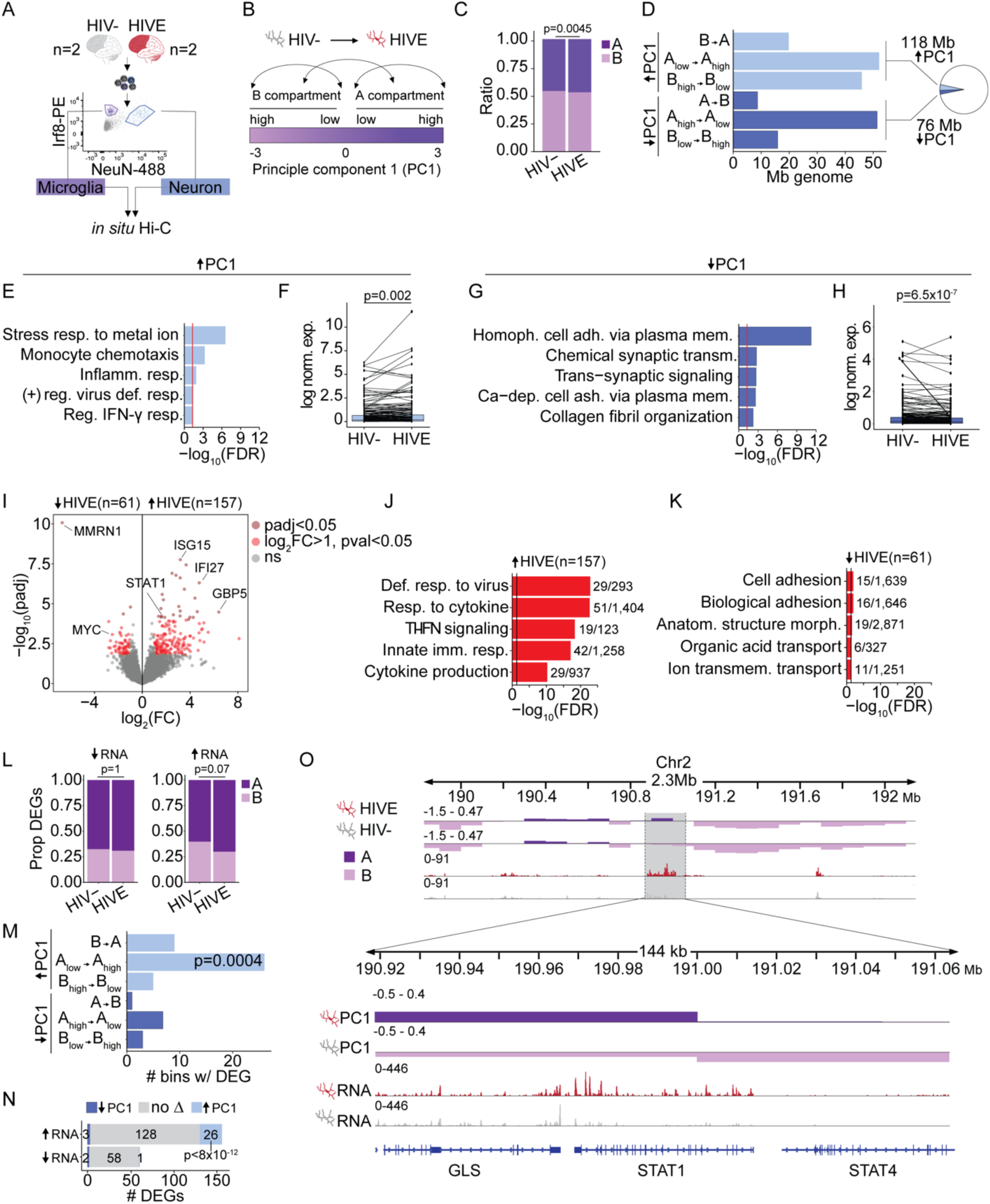
Chromosomal compartment remodeling and transcriptomic alterations in HIVE microglia. (**A**) Representative FANS plot for sorted NeuN and Irf8 immunostained neuronal and microglial nuclei which were used to generate *in situ* Hi-C libraries. (**B**) Schematic Hi-C compartment designation based on the eigenvector value of principle component 1 (PC1) of the Hi-C data. Arrows illustrate the different classes of compartment switches between HIVE and HIV-microglia. (**C**) Bar plot showing the proportion of the genome found in the A and B compartments, computed from Hi-C of HIV- and HIVE microglia. P-value reflects chi-square test. (**D**) Bar graph summarizing genome-wide compartment remodeling in HIVE microglia compared to HIV-microglia determined using dcHi-C. X-axis shows the number of Mb of the genome undergoing the compartment switch, Y-axis shows the different compartment switches from HIV-to HIVE with A-compartmentalization events in light blue (DPC1 HIVE-HIV->0, FDR<0.05) and B-compartmentalization events in dark blue (DPC1 HIVE-HIV-<0, FDR<0.05). Inset pie chart shows the total amount of significant A- and B-compartmentalization as a proportion of the genome. (**E,G**) Gene ontology enrichments and (**F,H**) differential expression for genes undergoing A-compartmentalization (E,F) and B-compartmentalization (G,H) in HIVE as compared to HIV-microglia. F,H) Paired line plot showing average log normalized expression of genes in compartments with a significant increase in PC1 in HIVE microglia. For visualization randomly selected subsets of 200 genes are shown. P-value reflects Wilcoxon signed rank test across all genes with expression levels above the 25^th^ percentile in either HIV- or HIVE microglia; (F) n=1030, (H) n=566. (**I**) Volcano plot showing snRNA-seq differential expression of the microglial cluster (Fig. 1C, Mg) from n=4 HIV- and n=8 HIVE brains. Points in dark red are genes with a padj<0.05. Points in lighter red are genes with a log2(FC)>1 and unadjusted pval<0.05. (**J-K**) Biological process GO enrichment for upregulated (J) and downregulated (K) genes. Black line marks significance for FDR=0.05, numbers next to each bar show the n genes differentially expression/the total n genes in that GO category. (**L**) Proportion of upregulated or downregulated microglial DEGs in A and B compartments in HIV- and HIVE microglia. P-value from chi-square test. (**M**) Bar plot with the number of 10kb bins undergoing compartment switching that contain a DEG shown on the x-axis. Fisher’s exact test compared to the proportion of the whole genome that underwent the indicated compartment switch. (**N**) Overlap between up- and downregulated DEGs in HIVE vs. HIV-microglia and all genes that underwent compartment switching, p-value from hypergeometric test, representation factor 5. (**O**) Representative browser shot of a 144kb window on chromosome 2 at the *STAT1/STAT4* gene locus showing Hi-C PC1 eigenvector value (top two tracks) with A (dark purple) and B (dark purple) compartment designations at a 10kb designation and normalized read counts from snRNA-seq Mg cluster (bottom two tracks) for HIV- (gray) and HIVE (red) brains.

To explore encephalitis-related changes to microglial genome structure, we first focused on megabase-sized (∼0.1-10Mb) chromosomal compartments, divided into gene-dense A-compartments with active gene expression and gene-poor B-compartments with low levels of active gene expression. Compartments are shaped by molecular forces linked to intranuclear phase separation (*26*), as compared to the self-folded topologically-associating domains (TADs) or loops, which primarily represent actively driven loop extrusion mechanisms in ensemble Hi-C data (*27*). Compartments were defined using the first principal component (PC1) eigenvectors at a 100kb resolution on a -3 (B) to +3 (A) scale (Fig. 2B). We observed a slight but significant shift in the proportion of the genome found in the A compartment in HIVE (47.3%) vs. HIV-microglia (46.2%, Chi-square p=0.0045, Fig. 2C). Analysis also revealed significant changes in A/B compartmentalization (detected by dcHi-C, FDR<0.05) encompassing 196Mb, or 6.4% of (haploid) genome in HIVE brain as compared to HIV-(Fig. 2D). This included 118Mb showing a significant increase in the PC1 eigenvector (HIVE > HIV-) corresponding to a switch to a more open conformation in HIVE microglia and 78Mb showing changes in the opposite direction, corresponding to compartment remodeling towards a more closed state (Fig. 2D, Data file S4). Importantly, functional pathway analysis revealed that genes positioned within genomic loci with significantly increased A-compartmentalization in HIVE (as compared to control) microglia showed significant enrichment for interferon (IFN) and other cytokine signaling pathways as well as myeloid chemotaxis and migration pathways (Fig. 2E, Data file S5-6). We then overlaid our Mg-specific A/B compartment maps with the transcriptomic maps constructed from the Mg cluster of our N = 7 HIVE and N=3 HIV-donors. The 1,940 genes within the regions of increased A compartmentalization in HIVE had significantly increased expression in HIVE as compared to HIV-microglia (Wilcoxon signed rank test, P = 6.5 x10^-7^) (Fig. 2F, see Data file S5 for all genes). This included a subgroup of N=335 genes that completely switched eigenvector direction from B to A, with significant enrichment in immune functions, including cytokine signaling, viral response, inflammatory pathways, and T-cell differentiation pathways (Fig. S4A). Genes that switched from Alow→Ahigh (N= 1,231) were enriched for metal ion stress response pathways due to a group of metallothionein genes, which have been shown to be involved in the immune response in general and in mediated monocyte resistance to HIV-induced apoptosis (*28, 29*). Genes that switched from Bhigh→Blow (N=308) were also enriched for immune functions, including lymphocyte chemotaxis, IFN, and interleukin pathways (Fig. S4B-C).

In striking contrast to these associations between A-compartmentalization and transcriptional upregulation of IFN and other cytokine and immune signaling genes in the HIVE microglia, gene ontologies of genomic loci (78Mb in total) undergoing repressive compartment remodeling, or B-compartmentalization, in HIVE microglia (PC1 HIVE < HIV^-^) were enriched for various homeostatic functions, such as cell adhesion, regulation of synaptic signaling, and neurodevelopment (Fig. 2G, Data file S6). This enrichment came primarily from Ahigh→Alow switching genes (N=687, Fig. S4D), while Blow→Bhigh genes (N=195) were enriched for ethanol and retinoic acid metabolism and A→B genes (N=145) had no significant GO enrichment (Fig. S4E). In agreement with the observed switch to a less active chromatin compartment, genes with increased B-compartmentalization were overall expressed at lower levels in HIVE microglia as compared to HIV-(Fig. 2H).

Next, in order to further test the link between compartment switching and transcriptional reprogramming in HIVE microglia, we inverted our order of analysis and first called N=218 differentially expressed genes (DEGs) in the Mg cluster of our snRNA-seq dataset (DESeq2, log2(Fold change) > 1, unadjusted P < 0.05) and then assigned their respective compartments (Fig. 2I, Data file S7). Top functional pathways for the N=157 upregulated genes, as with A-compartmentalized genes, were related to immune signaling, including the interferon (IFN) response with key transcription factors *STAT1* and *STAT2*, and IFN-stimulated genes *ISG15*, *IFI6*, *IFI27*, *IFI44*, and *IFIH1* (Fig. 2J, Data file S6). In addition, upregulated expression with A-compartmentalization was found of several members of the guanylate binding proteins (GBPs), *GBP1*, *2*, *4*, and *5*, which interact with the inflammasome to regulate cytokine production and pyroptosis (Fig. 2J) (*30*). We also observed overlap of functional pathways for the N=61 downregulated genes and B-compartmentalized genes, with both showing enrichment of cell adhesion genes (Fig. 2K, Data file S6).

Of note, from the 157 DEGs upregulated in HIVE Mg, 70% were found in the A compartment in HIVE microglia as compared to 60% in HIV-microglia (Chi-square p=0.07, Fig. 2L). Furthermore, there was significant enrichment for Alow→Ahigh compartment switching among DEGs (Fig. 2M). This enrichment came from upregulated DEGs, of which 17% were found in regions with increased A-compartmentalization in HIVE, representing a significant, 5-fold enrichment over chance (hypergeometric p<8x10-12, Fig. 2N). A representative example of a genomic locus undergoing A-compartmentalization in conjunction with transcriptional upregulation at the *STAT1/STAT4* locus, is shown in Figure 2O. In contrast to upregulated DEGs, we did not observe significant changes in compartment location of downregulated DEGs, suggesting other mechanisms may be involved in transcriptional regulation of these genes (Fig. 2L, N). Furthermore, there was a significant linear correlation (Pearson R=0.41, p=1x10^-11^) of the log2(fold change) in expression of the 218 DEGs in HIVE and HIV+ as compared to HIV-, suggesting the same changes may be present to a lesser extent in HIV+ brain without high levels of viral expression (Fig. S4F). Together, these results suggest that HIV induces open chromatin configuration in microglia that accompanies transcriptomic activation of inflammatory pathways. The 3D remodeling of microglial genomes in the HIVE brain was not limited to compartment architectures. Chromosomal loops, defined as distinct contacts between two loci in the absence of similar interactions in the surrounding sequences (*31*), showed large-scale remodeling genome-wide between groups, with N= 5,959 loops specific to HIVE, N=2,799 loops specific to HIV^-^, and N=1,493 loops shared between HIVE and HIV^-^ microglia (Data file S8). Genes found only within the span of HIVE loops were enriched for functions including STAT signaling, cell adhesion, and nervous system development, which were similar to pathways we observed enriched in compartment and differential expression analyses (Fig. 3A, Data file S6). Furthermore, microglial DEGs were significantly enriched in HIVE-specific loops, while no enrichment was observed for HIV-loops (Fig. 3B). A representative example is shown in Figure 3C, with HIVE-specific loop formation over the differentially expressed oligoadenylate synthetase (OAS) cluster genes on chromosome 12, encompassing *OAS1*, *2*, and *3*, which showed upregulated expression in HIVE microglia and are defined as IFN-stimulated genes that encode RNases capable of cleaving viral RNAs (*32*). We observed specific loops in HIVE microglia that spanned from a region upstream of the OAS cluster to enhancers distal to the cluster, while loops anchored in the flanking sequences of the OAS cluster were common both to HIVE and HIV^-^ microglia.

**Figure 3.**
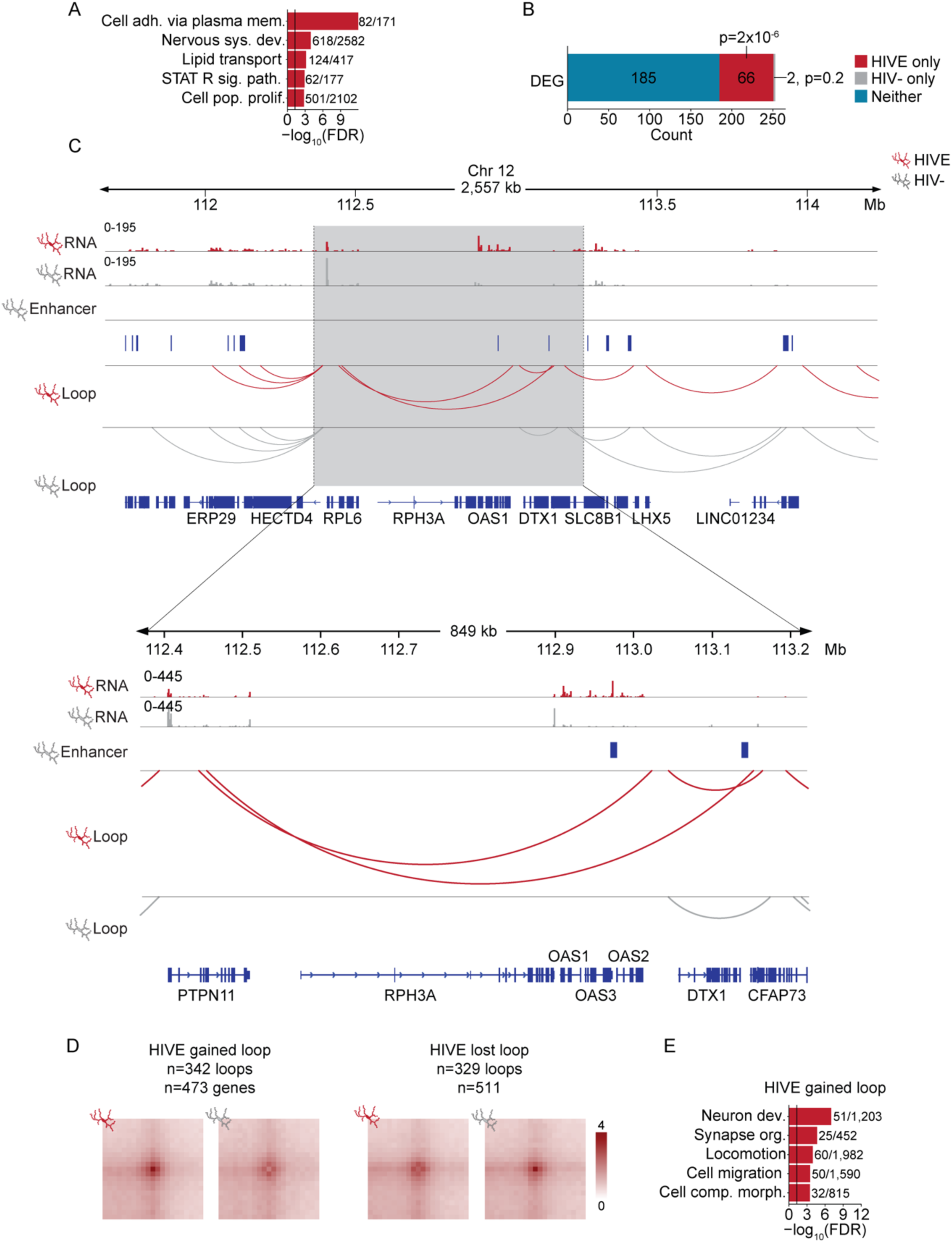
Chromosomal loop remodeling and transcriptomic alterations in HIVE microglia. (**A**) Gene ontology enrichment for genes found in HIVE chromosomal loops. Black line indicates significance threshold at FDR=0.05. Numbers indicate the number of genes in the dataset/the number of genes in the GO category. GO categories: cell adh. via plasma mem., cell adhesion via plasma membrane; nervous sys. dev.; nervous system development; STAT R sig. path., STAT receptor signaling pathway; cell pop. prolif., cell population proliferation. (**B**) Bar graph showing the number of DEGs that are found only in HIVE loops, only in HIV-loops, or in neither HIVE or HIV-specific loops. Hypergeometric test for the overlap between the DEGs and genes in HIV- or HIVE-specific loops using all genes as a background. (**C**) Representative locus showing HIVE-specific loops associated with differential expression of the OAS genes on chromosome 12. Normalized read counts from snRNA-seq Mg cluster (top two tracks) for HIV- (gray) and HIVE (red) brains, enhancers called from microglial H3K27ac ChIP-seq data (middle track), and Hi-C loops called by Mustache (bottom two tracks) for HIV- (gray) and HIVE (red) are shown. (**D**) Contact frequency maps shown for examples of differential loops that were called on pooled loops from HIVE, HIV-, and combined HIVE and HIV-. For all loops, normalized contact frequencies in HIVE and HIV-queried using a linear regression model and loops with p<0.05 were considered significant. Numbers of loops gained and lost in HIVE and the number of genes located in the anchors of gained/lost loops are indicated. (**E**) Gene ontology enrichment for genes found in HIVE gained loops. Black line indicates significance threshold at FDR=0.05. Numbers indicate the number of genes in the dataset/the number of genes in the GO category. GO categories: Neuron dev., neuron development; synapse org., synapse organization; cell comp. morph., cellular component morphogenesis.

Loop numbers are highly dependent on Hi-C read depth. Thus, while the enrichment observed for DEGs in HIVE-specific loops controlled for the differences in loop number between HIVE and HIV-microglia, we performed further loop analysis controlling for read depth. We identified 342 loops that were gained in HIVE and 329 loops that were lost in HIVE (Fig. 3D, Data file S8). HIVE gained loop anchors contained 473 genes (Data file S9), which again were enriched for genes involved in pathways similar to those observed in compartment and differential expression analysis, including synapse organization and cell migration (Fig. 3E, Data file S6). Loops that were lost in HIVE contained 511 genes (Data file S9), which, as with the initial loop analysis, no significant GO enrichment.

### Subhead 3: Neuronal and non-neuronal integration site mapping in the HIV infected brain

Our single nuclei transcriptome profiling from cortical gray and adjacent subcortical white matter showed that HIV RNA is detectable in a subset of HIVE microglia, with a smaller contribution of other, overwhelmingly non-neuronal, cell types (Fig. 1E-F). Therefore, for genome-wide integration site sequencing (IS-seq), we sorted and separated neuronal (NeuN+) and non-neuronal (NeuN-) nuclei from FC/WM by FANS (Fig. 4A) followed by construction of cell-type specific integration site (IS) sequencing (IS-seq) libraries (Fig. S5A-B). To validate our IS-seq methods, we generated IS-seq libraries from four JLat lymphocyte lines each harboring a single, previously published IS, and we identified the correct IS in each case with no spurious IS detected (Fig. 4A, Fig. S5C). We further generated IS-seq libraries from sequential dilutions of the JLat clones with uninfected Jurkat DNA, which revealed an overall 1.6% sensitivity of IS detection, a number that is 6-fold higher than previous studies using similar methodology (Fig. S5D) (*33*). Importantly, DNA FISH labeling of nuclei from HIVE brain with an HIV-specific 4kb probe showed a single integrated provirus in each infected nucleus, suggesting that surviving, infected cells in the brain are not targeted for repeated infection, similar to what has been observed in peripheral lymphocytes (Fig. 4A) (*34, 35*). Therefore, to get better insight into the genomic positioning of viral insertions in brain cells, we generated 53 cell-type specific IS-seq libraries from FC/WM sorted neuronal and non-neuronal nuclei from 25 unique donors (n=7 HIVE, n=18 HIV+; Table S1). 20 HIV-control IS-seq libraries were sequenced, 8 from uninfected Jurkat DNA and 12 from HIV-FC/WM sorted nuclei from 6 unique donors. An additional 20 libraries were prepared from 14 HIV+, non-encephalitic donors and 8 libraries from HIV-samples that failed to produce sufficient library due to low or absent viral DNA (Table S1). From the sequenced libraries, we identified n=1,254 IS, the overwhelming majority (n=1,221/1254, 97%) of which came from NeuN-nuclei (Data file S10). Of note, IS were detected in 85% (6/7) of HIVE FC/WM donors and 28% (5/18) HIV^+^ donors, including 2 subjects diagnosed with CD8+ T-cell encephalitis, an unusual inflammatory brain disorder often seen in cART-medicated individuals, but also in treatment-naïve individuals (*36*), and 1 elite controller (Fig. 4B).

**Figure 4.**
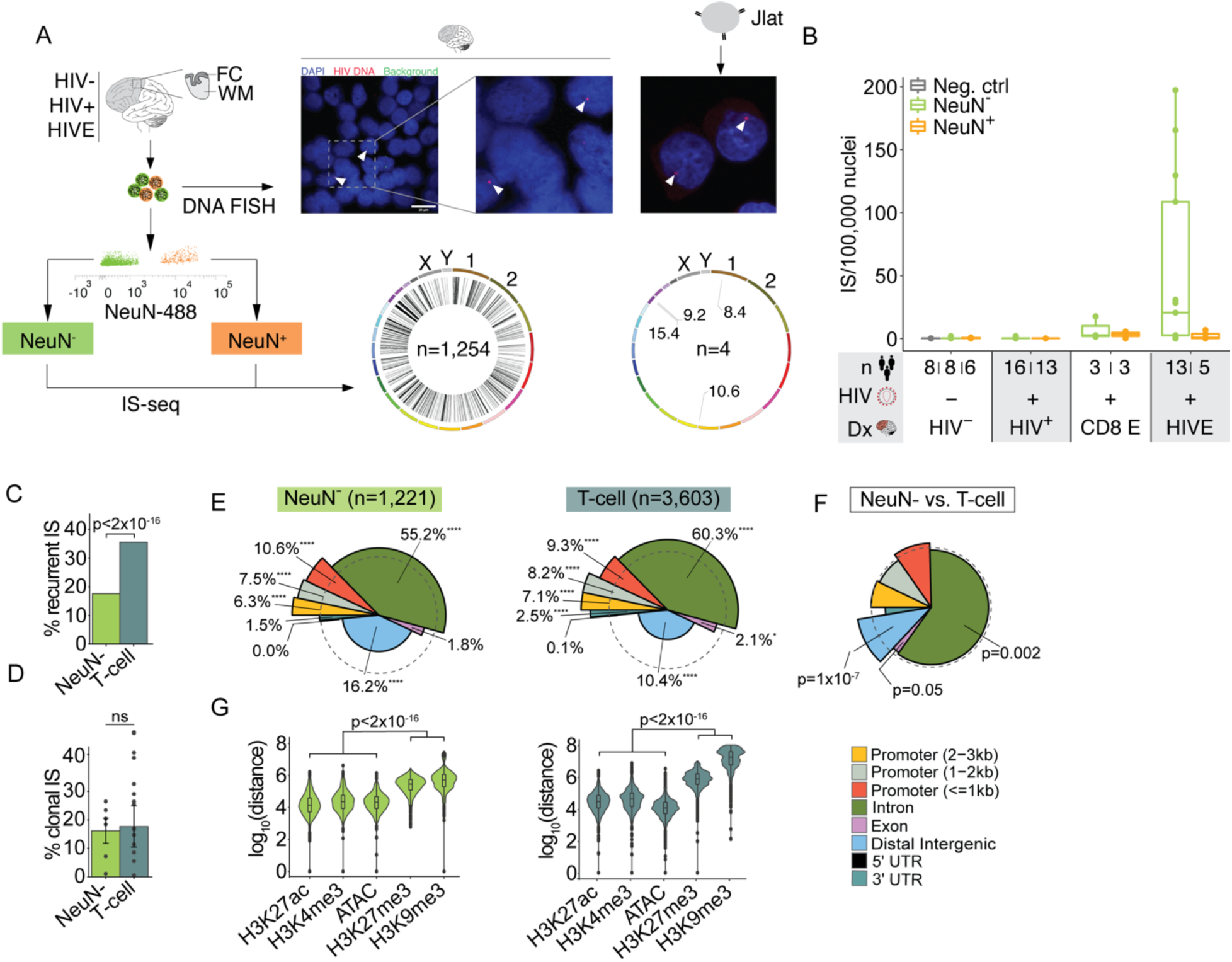
Genomic and epigenomic features of IS in human brain. (**A**) Schematic showing a representative FANS plot for sorting of neuronal and non-neuronal nuclei for IS-seq or FC/WM and JLat cells for comparison. Representative images are shown of DNA FISH from HIVE nuclei (left) and JLat cells (right). Arrowheads indicate positive HIV DNA FISH signal. Genome-wide IS shown on circos plots for brain (left, n=1,254 IS) and for 4 JLat clones (right, n=4 IS). (**B**) Boxplot of the number of IS detected per 100,000 nuclei (y-axis) shown for negative control DNA from uninfected Jurkat cells (Neg. Ctrl., gray), NeuN-(green), and NeuN+ (orange) nuclei. Samples are grouped by diagnosis (Dx) on the x-axis into HIV-, HIV+, CD8 encephalitis (CD8 E), and HIVE. (**C-D**) Percent of IS that are recurrent (C) or clonal (D) for all NeuN-and published in vivo T-cell IS (T-cell) (*37–42*). Significance testing by chi-square (C) or two-way ANOVA (D). (**E**) Spie charts comparing IS distributions across 8 genomic features as indicated to overall feature distributions in the genome. The angle of the pie slice represents the proportion of genome occupied by that feature, while the height of the pie slice represents the relative feature proportion in the IS dataset as compared to the genome, dashed circle demarcates equal proportions and % denote the IS feature percent. Chi-square test. * 0.01 < p < 0.05, ** 0.001 < p < 0.01, *** 0.0001 < p < 0.001, **** p<0.0001. (**F**) Spie chart comparing NeuN-IS distribution to T-cell. Chi-square test. G) Violin plots showing the distance between IS and the nearest peak for open chromatin markers H3K27ac, H3K4me3, and ATAC-seq and repressive chromatin marks H3K27me and H3K9me3. NeuN-IS are compared to NeuN-specific ChIP- and ATAC-seq and T-cell IS are compared to Jurkat ChIP- and ATAC-seq. Kruskall Wallis rank sum tests.

We first compared the integration map from our NeuN^-^ population of nuclei with three independent T-cell datasets: 1) n=3,603 IS from patient T-cells infected *in vivo* that were previously published (*37–42*) n=687 IS from peripheral blood mononuclear cells (PMBCs) for which raw sequencing files were available and processed by our pipeline (see Methods, Data file S10) (*41*), 3) n=52,190 IS we sequenced from Jurkat cells infected *in vitro* (Data file S10). Of note, the IS-seq map from NeuN-nuclei showed a significantly lower rate of recurrent integration events, defined as >1 separate integration event into the same gene (Fig. 4C). This effect was consistent when compared to each of the 3 viral integration maps from peripheral lymphocytes (Fig. 4C, Figure S6A). However, clonal integration in non-neuronal nuclei (16% average across donors) was of similar magnitude compared to *in vivo* T-lymphocyte (18%) and PMBC (12%) datasets (Fig. 4D, Fig. S6B), while clonal events in the Jurkat cells, harvested 48h post-infection, were significantly lower (0.6%) (Fig. S6B). Thus, it appears infected microglia proliferate during HIVE.

Furthermore, IS genomic annotations revealed, both for NeuN^-^ HIVE and peripheral T-cells, significant enrichments for promoter proximal sequences, introns, and 3’UTRs (Fig. 4E, Figure S6C). However, when we compared NeuN-IS to *in vivo* T-cell IS, we observed significantly more integration into distal intergenic regions and significantly less integration into both introns and exons in NeuN-cells (Fig. 4F). Next, we explored local epigenomic features at viral insertion sites by studying T-cell IS in relation to Jurkat publicly available ChIP-seq datasets and NeuN-IS in relation to publicly available or newly generated (see Methods and Data file S1) NeuN-ChIP-seq datasets for four types of histone H3 covalent modifications differentiating between transcription-facilitative (H3K4me3 trimethyl-lysine 4 and H3K27ac acetyl-lysine 27) and repressive (H3K9me3 trimethyl-lysine 9 and H3K27me3 lysine 27) nucleosomal organization, together with open chromatin profiles by ATAC (Assay for Transposase Accessible Chromatin)-seq. Indeed, in both NeuN-cells and peripheral lymphocytes, IS were located ∼10^4^ bp from the nearest open/facultative chromatin mark but were 10^6^-10^8^bp from the nearest repressive marks (Fig. 4G, Fig. S6D).

Interestingly, we observed that *LEDGF* and *CPSF6*, the two cellular factors involved in mediating integration site selection in T-cells, were expressed at significantly lower levels in microglia as compared to all other cell clusters, including the lymphoid cluster, in our snRNA-seq data (Fig. S7A-B). We leveraged publicly available ChIP-seq data from U2OS cells (Data file S1), and observed that while there was enrichment for CPSF6 binding at T-cell IS, there was no enrichment at NeuN-IS (Fig. S7C-D). A recent work published the first *in vitro* microglial IS in the C20 microglial cell line and did not observe the same differences in IS patterns as we observed here compared to T-cells (*43*). However, based on published RNA-seq data, expression of *LEDGF* and *CPSF6* are significantly higher in C20 cells compared to primary microglia (Fig. S7E-F). Both in our snRNA-seq data and published RNA-seq data (*44*), the discrepancy in expression appears to be greatest for *LEGDF*. Thus, while general IS patterns in the brain are similar to T-cells, we observe differences in the rates of recurrent integration, integration into intergenic regions, and expression and binding of integration targeting host factors.

### Subhead 4: Viral integration is associated with microglial transcriptional and 3D-genomic features

We next studied the relationship between viral integration and transcription. We observed that genes containing IS (IS genes) were expressed at significantly higher levels in microglia as compared to non-immune cell types in the brain but were expressed at similar levels in Mg, ImOl, and Lymph clusters (Fig. 5A). This is in line with our snRNA-seq data, which showed viral transcription predominantly in the Mg and ImOl clusters (Fig. 1E-F). Similarly, microglia showed highest level of expression for IS genes most variably expressed across our eight principal cell types (Fig. 5B).

**Figure 5.**
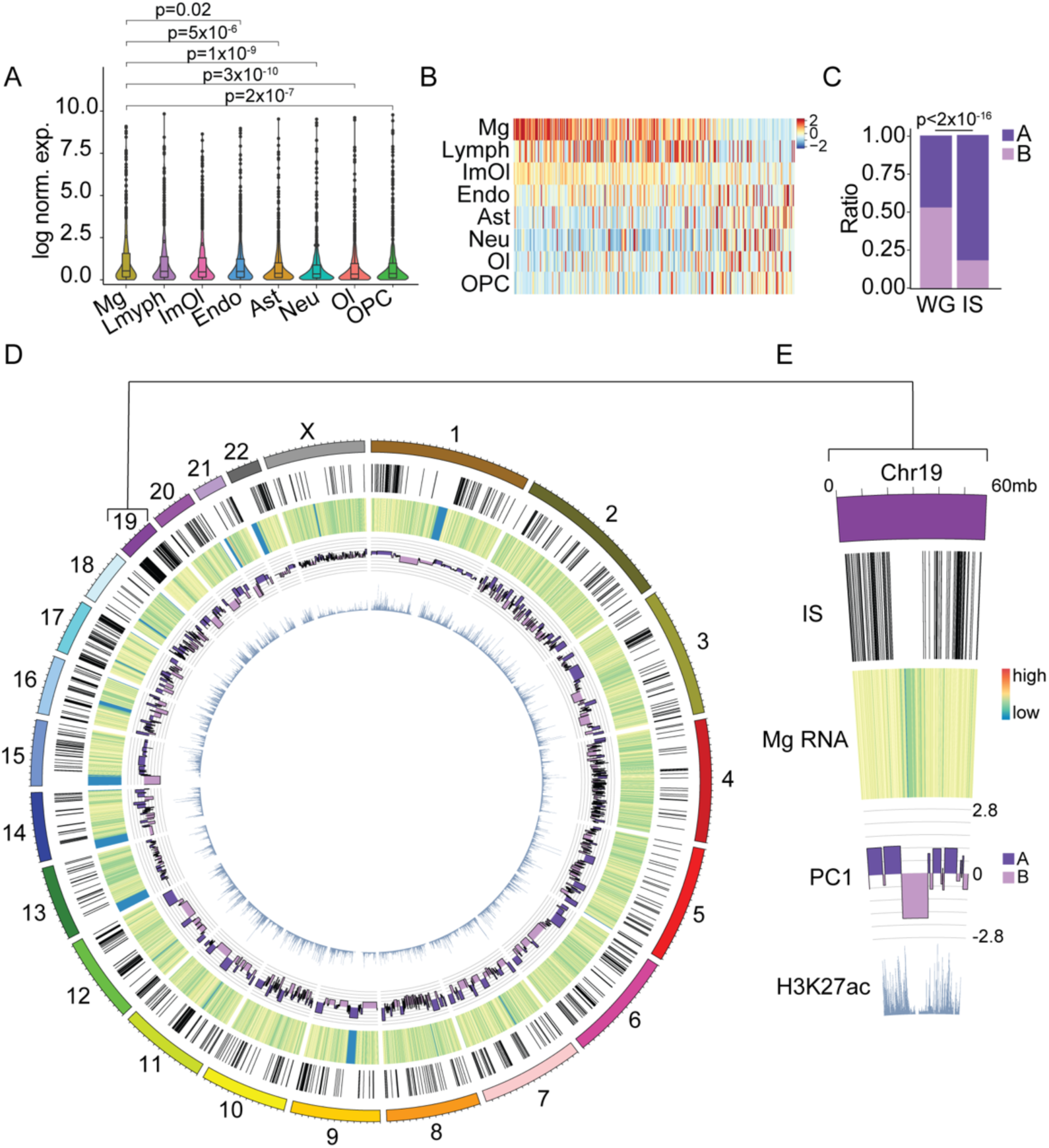
Viral integration is associated with microglial transcription and genome structure. (A) Violin plot based on snRNA-seq FC/WM data showing cluster-specific expression of all genes containing IS. For cell types composed of more than 1 subcluster, values were averaged across all cells. Wilcoxon rank sum test of Mg expression vs. other cell types. (**B**) Heatmap showing scaled expression of the top 300 genes most variably expressed across cell types that contain IS. (**C**) Proportion of the whole genome (WG) and NeuN-IS found in HIVE microglial A and B compartments. Chi-square test. (**D-E**) Genome-wide (D) or chromosome 19 specific (E) circos plot displaying from the outside in NeuN-IS, heatmap of log scaled HIVE Mg snRNA-seq read counts, HIVE Mg Hi-C A/B compartment designation expressed as PC1 value, and NeuN-H3K27ac ChIP-seq reads.

Consistent with viral integration preference at sites of open chromatin marked by activating histone modifications, the NeuN-brain IS were overwhelmingly located in the Hi-C defined A-compartment (82%, Fig. 5C). These patterns of integration into the A compartment and regions of active gene expression could be observed genome-wide (Fig. 5D). Chromosome 19, which is the chromosome with the highest gene density and integration site density, demonstrated global IS patterns especially clearly (Fig. 5E). IS were sharply confined to A-compartment rich locations in the ∼20Mb spanning flanking sites of the chromosome, with complete sparing of repressive B-compartment chromatin encompassing ∼20Mb in the chromosomal center.

### Subhead 5: Viral integration is linked to activation-related reorganization of the microglial genome

In T-cells, 3D genome reorganization in *in vitro* models of immune activation has been shown to impact integration site selection (*45*). We wanted to see if this was also the case *in vivo* in microglia. We observed a significant increase in the proportion of IS in the A compartment in HIVE as compared to HIV^-^ microglia (82% vs. 78%, Chi-square p=0.02; Fig. 6A). We then looked to see whether IS were found in regions of significant compartment switching. 7.6% of IS were found in regions of compartment switching (Fig. 6B), with strong preference for regions undergoing significant A-compartmentalization (Chi-square, p=0.06) (Fig. 6B). Finally, we observed that IS were associated with a significantly lower insulation score, a measure of domain insulation with lower scores defined as being more insulated, in HIVE as compared to HIV-microglia (Fig. 6C), while the whole genome had a significant increase in insulation or a weakening of domain structure (Fig. S8A). An example of this is shown for 2 IS on chromosome 11, where strengthened intra-domain interactions drive a drop in the insulation score at the site of integration (Fig. 6D). Thus, HIV preferentially integrates in regions with increased intra-domain interactions in HIVE microglia.

**Figure 6.**
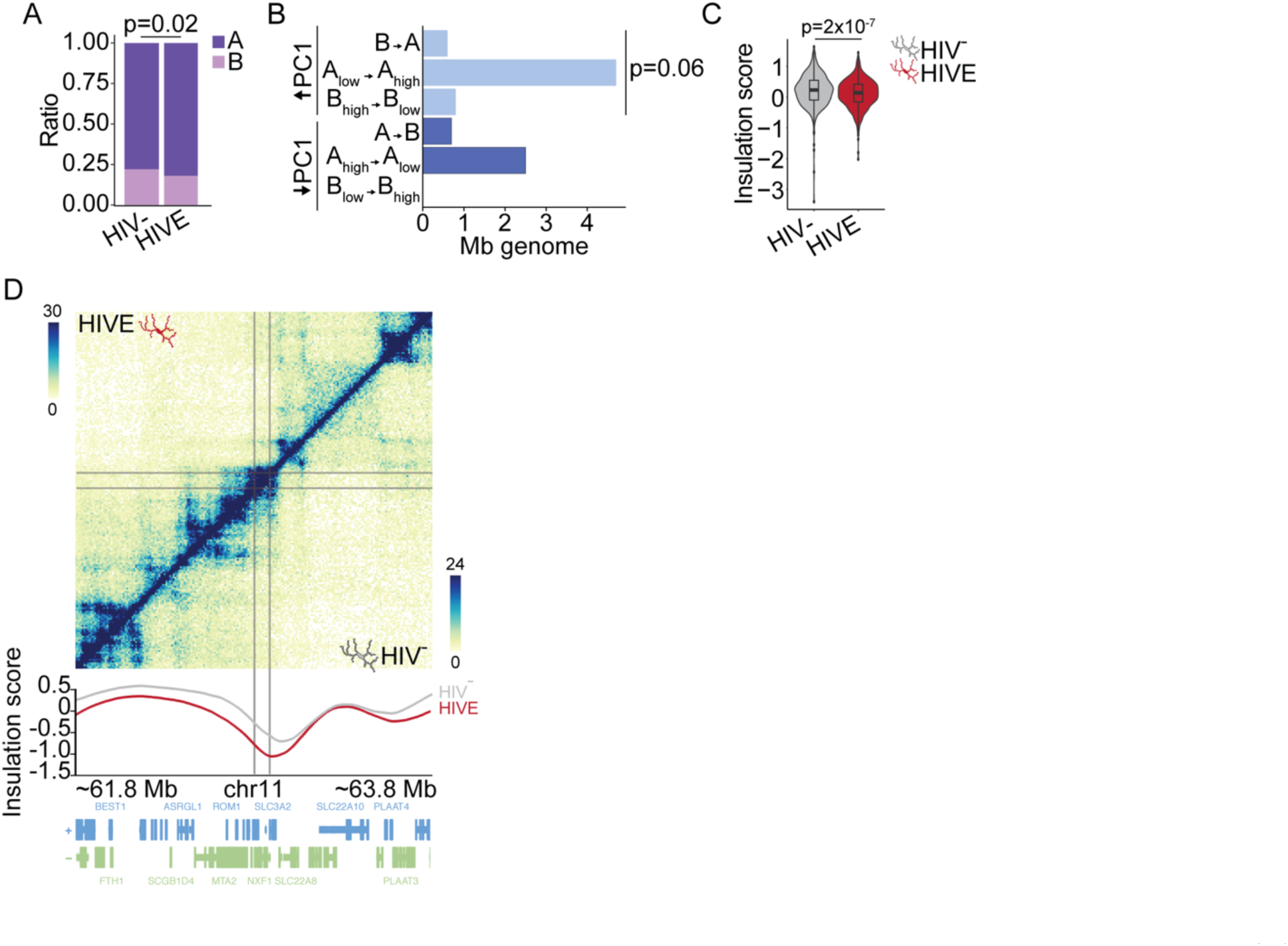
Viral integration is associated with 3D-genomic restructuring in HIVE. (**A**) Proportion of NeuN-IS found in A and B compartments in HIV- and HIVE Mg. Chi-square test. (**B**) Bar graph summarizing compartment remodeling at IS in HIVE microglia compared to HIV-microglia determined using dcHi-C. X-axis shows the number of Mb of the genome in bins that have an IS and a significant change in PC1, Y-axis shows the different compartment switches from HIV- to HIVE with A-compartmentalization events in light blue (DPC1 HIVE-HIV- >0, FDR<0.05) and B-compartmentalization events in dark blue (DPC1 HIVE-HIV- <0, FDR<0.05). (**C**) Violin plot showing insulation score at IS in HIV- and HIVE microglia. Wilcoxon rank sum test. (**D**) Hi-C contact frequency maps of HIVE (top half) and HIV- (bottom half) microglia shown for representative integration sites (gray lines) on chromosome 11. Insulation score is plotted below.

Regions with the greatest changes in insulation, both genome-wide and specifically for those regions containing IS, were significantly enriched for YY1 and CTCF binding from publicly available ChIP-seq data (Fig. S8B-C). YY1 and CTCF are both chromatin structuring proteins that mediate either enhancer-promoter or insulating loops respectively (*46, 47*). This suggests that YY1 and CTCF binding may play a role in chromatin restructuring in HIVE.

### Subhead 6: Viral transcription is linked to microglial activation

Integration is a critical step in the HIV life-cycle because it allows for efficient transcription of viral genes (*48, 49*). Thus, we next studied viral transcription using our snRNA-seq data and were surprised to see that HIV was the second most highly expressed transcript out of >17,000 expressed transcripts in HIV RNA+ microglial nuclei (Fig. 7A). This suggests that HIV takes over the cellular transcriptional landscape. Indeed, we observed that HIV RNA+ nuclei tended to cluster together when we performed sub-clustering of HIVE microglia, even though HIV transcripts were excluded when performing the sub-clustering (Fig. 7B). Furthermore, when performing principal component analysis of HIV RNA+ and RNA-microglia from each of our 5 HIVE donor samples in which we detected viral transcription, the HIV RNA+ microglia showed a highly consistent shift along PC1 and PC2 (Fig. S9A), indicating a consistent effect of active viral transcription on cellular gene expression across these different subjects. Differential expression analysis of HIV RNA+ vs. RNA-Mg revealed 69 downregulated genes, which were enriched in pathways related to cell adhesion and neuronal function (Fig. 7C-D). Over half of these genes are the targets of CTBP2 and SUZ12, members of the polycomb repressive complex 2 (PRC2) which were both expressed more highly in HIV RNA+ microglia, suggesting that PRC2 may play a role in repressing transcription of genes in HIV-infected cells (Fig. S9B-C). There were 20 significantly upregulated genes in HIV RNA+ microglial nuclei that were enriched for phagocytic and cell migratory functions (Fig. 7C, E). Importantly, very similar gene ontologies and functional pathways emerged in our snRNA-seq and Hi-C analyses comparing HIVE microglia as a whole to microglia from HIV-brain (described in Fig. 2). Therefore, we wanted to further compare HIV RNA+ microglia with the overall population of HIVE microglia. We observed a strong, highly significant positive linear relationship between the log2(Fold Change) values of RNA+ vs. RNA-Mg and all HIVE Mg vs. HIV-Mg (Fig. 7F), suggesting that viral transcription occurs in activated microglia that drive differential expression in HIVE. In line with this, for the important immune related genes, *HLA-B* and *STAT1*, we observed a gradient in expression that was lowest in HIV-microglia, intermediate in HIVE microglial nuclei not harboring HIV transcripts, and highest in HIV RNA+ HIVE microglia, (Fig. 7G). HIVE HIV RNA-microglia still showed evidence of activation, as there was a high degree of correlation between HIVE HIV RNA- and RNA+ differential expression as compared to HIV-microglia (Fig. S9D). However, the magnitude of the fold change in expression of DEGs was higher for HIV RNA+ as compared to RNA-microglia, further reinforcing that HIV RNA+ Mg are the most activated (Fig. S9E). Taken together, these findings suggest that viral transcription detectable on the level of the single nucleus is linked to a subset of highly activated microglia.

**Figure 7.**
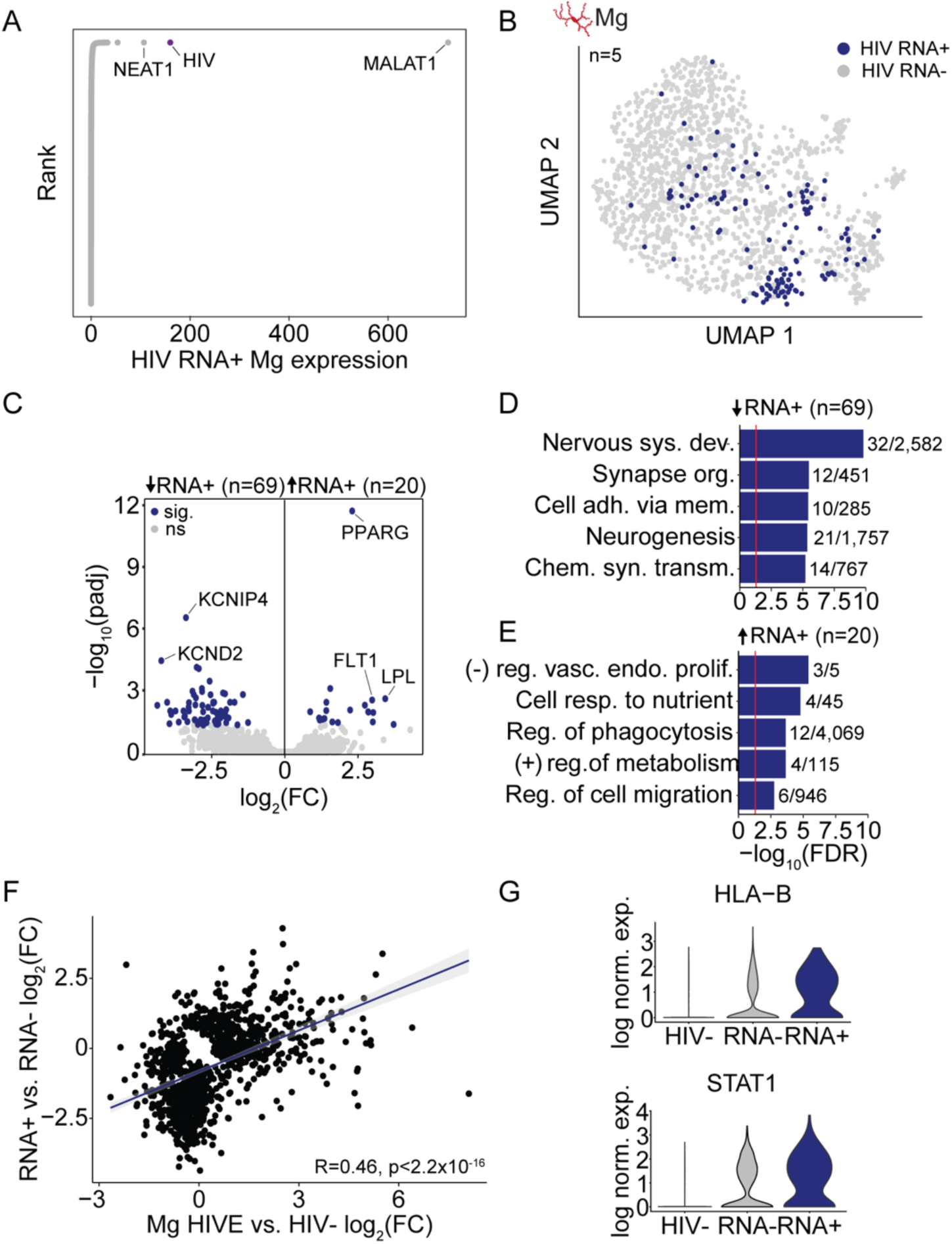
Viral transcription is associated with microglial activation. (**A**) Rank plot showing the log normalized expression of each gene in HIV RNA-expressing (RNA+) HIVE microglia ranked by level of expression. (**B**) HIV RNA expression mapped onto sub-clustered microglia from HIVE brains that had viral transcription present (n=5). Cells with HIV transcripts detected (HIV RNA+) are shown in blue, those without (HIV RNA-) are shown in gray. (**C**) Volcano plot showing log2(fold change) and -log10(adjusted p-value) for each gene. Points in blue are genes that are significant with a padj<0.05. (**D-E**) Biological process GO enrichment for downregulated (D) and upregulated (E) genes. Red line marks significance for FDR=0.05, numbers show the number of genes for each GO category that are differentially expressed/the total number of genes in that GO category. GO categories: Nervous sys. dev., nervous system development; cell adh. via plasma mem., cell adhesion via plasma membrane; chem. synaptic transm., chemical synaptic transmission; (-) reg. of vasc. endo. prolif., negative regulation of vascular endothelial proliferation; cell resp. to nutrient, cellular response to nutrient; (+) reg. of metabolic process, positive regulation of metabolic process. (**F**) Pearson correlation of log2(fold change) in gene expression between RNA+ vs. RNA-HIVE microglia (Y-axis) and all HIVE microglia vs. HIV-microglia (X-axis) for genes with an unadjusted pval<0.05 in at least one comparison. (**G**) Violin plots showing log normalized expression of HLA-B and STAT1 in HIV-microglia, HIVE HIV RNA-microglia, and HIVE HIV RNA+ microglia.

## DISCUSSION

Deeper understanding of the invasion and spread of HIV in the brain and its clinical sequelae, including HAND, requires insight into the epigenomic and transcriptomic context of the retroviral infection in the human brain. Here, we built an integrative dataset from 77 sequencing files, including snRNA-seq, Hi-C, IS-seq, and ChIP-seq, providing a critically needed neurogenomics resource from postmortem brain of infected donors and controls. By providing genome-scale maps for viral integration sites and their disease-associated chromosomal conformation and single nuclei expression states, our resource greatly extends previous, tissue-homogenate based gene expression profiling and integration site analyses in cell culture models (*43*).

While HIVE is no longer common in the cART era, it is useful as a model to understand how active, unrestricted viral infection impacts the brain, as all individuals undergo a period of unrestricted viral replication prior to initiating cART (*50, 51*). In support of this, patients occasionally have neurologic symptoms during acute infection, so it is possible the brain at this stage may resemble more of a HIVE-like state (*52*). It will be interesting to utilize simian immunodeficiency virus (SIV) models (*53*) or humanized mouse models (*54*) to assess whether the neurogenomics of acute infection resonates with some of the findings reported here for the HIVE postmortem brain. Moreover, HIVE remains a significant concern for an estimated 9 million PLWH worldwide believed not to be on cART (*55*), given that in the pre-cART era approximately one quarter of patients develop HIVE in the setting of severe immunosuppression (*56, 57*).

In any case, our single nuclei profiling, and earlier work in HIVE bulk tissue (*58, 59*), all point to elevated expression of interferon (IFN) response and complement cascade genes, and decreased expression of genes related to bioenergetics and neuronal function, indicating an adaptive shift in the microglia of the infected brain from homeostatic, neuronal support functions to inflammation. This may contribute to the observed neuronal death in HIV infection in the absence of direct neuronal infection (*60*). Importantly, according to our microglia-specific Hi-C data from HIVE and non-infected controls, chromosomal conformations at the site of differentially expressed genes undergo a profound reorganization, with altogether 182 megabases of microglial genome—in total length roughly equivalent to the size of human chromosome 5—undergoing a change in A/B compartment structures, together with rewiring of thousands of contact-specific loops and a genome-wide weakening of the kilo-to megabase-scaling TAD domains. Specifically, sites of up-regulated gene expression, including many genes involved in interferon response and viral defense pathways, undergo a reconfiguration to a transcriptionally more permissive and open chromatin environment.

Critically, these activation-associated changes to microglia appear to potentiate viral infection, both in terms of integration and viral transcription. We observed a strong preference for integration into open chromatin, defined in terms of epigenomic and 3D-genomic signatures, similar to what has been previously reported in T-cells (*37, 39, 45, 61*). Integration was enriched in areas of 3D remodeling in HIVE, which may represent a feed-forward cycle of integration and activation that further propagates viral spread. We also observed that viral transcription occurred in a subset of highly activated microglia, where it essentially highjacked the cell to become the second most highly expressed transcript. This underscores the capability of HIV for extremely productive, and ultimately detrimental, seeding and spread in the brain’s susceptible cell population. The presence of clonal integration sites further points to the ability of infected cells to replicate and expand the pool of CNS virus.

Importantly, we also observed lower rates of recurrent integration and higher rates of integration into intergenic regions in the brain as compared to T-cells, which may be caused by lower levels of expression of the integration targeting factors, *LEDGF* and *CPSF6*. In macrophages, lower rates of genic integration have also been reported and suggested to be due to lower expression of *LEDGF* (*62*). These findings highlight the need to further study integration in microglia, and our analysis showing certain microglial cell lines express *LEDGF* and *CPSF6* at much higher levels means that this work must be done in primary or potentially induced pluripotent stem cell-derived microglia. Differences in integration site targeting have implications for HIV cure strategies as latency reversing agents used in shock-and-kill approaches have different efficacy depending on the integration site (*63*).

Our study also suggests that there are similar, lower magnitude transcriptional changes present in HIV+, non-encephalitic brain. This is important in terms of understanding the molecular mechanisms of HAND in the setting of viral suppression. However, given the smaller sample size analyzed for HIV+ tissue by snRNA-seq, these findings require follow up study with larger numbers of individuals. The datasets presented here will provide a rich resource to better understand cell-type, sequence- and chromatin status-specific mechanisms that govern HIV integration, expression and eventually, silencing in the human brain.

## MATERIALS AND METHODS

### Study design

Utilizing frozen postmortem brain tissue from the Manhattan HIV Brain Bank, we performed cell-type-specific, genome-wide sequencing of three different types: 1) single-nucleus RNA-sequencing (snRNA-seq), 2) *in situ* Hi-C, and 3) HIV integration site sequencing (IS-seq). We assayed tissue from three different donor types: 1) HIV-uninfected control (HIV-), HIV-infected (HIV+), and HIV-infected with HIV encephalitis (HIVE), who served as a positive control for active CNS viral infection. The diagnosis of HIVE was made by a board-certified neuropathologist on routine histopathology, based on the presence of a microglial nodule encephalitis with characteristic multinucleated syncytial cells and staining for HIV p24 antigen. Other HIV-infected individuals (the HIV+ group) did not have this histologic evidence of active HIV brain replication, and were variable with regard to cART status and virologic control prior to death. Donor characteristics, including ante-mortem plasma HIV loads, CD4 counts, and cART status, are further described in Table S1. There were a small number of donors for whom some of this information was not available.

Stringent quality controls were used for each assay performed and are described in depth in their respective sections below. Briefly, for snRNA-seq we applied filters to remove low quality nuclei from analysis and performed experiments in which mouse tissue was mixed with human tissue to determine a reliable HIV read count threshold. For Hi-C, we used paired HIV-and HIVE samples and processed all samples in parallel, at the same time. Correlation analysis was used to ensure biological replicates were aligned. For IS-seq, each time the assay was run, a positive and negative control were included. This allowed us to ensure a lack of contamination in IS-seq libraries and that, when a particular sample failed to yield library, this was due to low levels of HIV rather than a failure of the assay.

### Human tissue

Human autopsy brain samples were collected by the Manhattan HIV Brain Bank (MHBB, U24MH100931), using protocols under the supervision of the Icahn School of Medicine at Mount Sinai (ISMMS) Institutional Review Board. Written informed consent was obtained either from decedents or their primary next of kin. Brain processing and histologic assessment were completed as previously described (*64*).

### Nuclei isolation

Nuclei were isolated from 200-300mg of human postmortem frontal lobe tissue (cortex and underlying white matter) as previously described (*65*). Briefly, tissue was homogenized in a hypotonic lysis buffer (0.32M sucrose, 5mM CaCl2, 3mM Mg(Ace)2, 0.1mM EDTA, 10mM Tris-HCl pH 8, 1mM DTT, 0.1% Triton X-100). A sucrose gradient (1.8M sucrose, 3mM Mg(Ace)2, 1mM DTT, 10mM Tris-HCl pH8) was layered under the homogenate and nuclei were pelleted via ultracentrifugation at 24,000rpm for 60 minutes at 4°C.

For snRNA-seq nuclei were resuspended in 1mL 0.04% BSA in DPBS. In cases where mouse and human tissue were mixed, cortical tissue from wild-type C57BL/6 mice was combined with human tissue prior to homogenization.

For Hi-C, Tissue was homogenized as described above, and tissue homogenate was fixed in 1% formaldehyde at room temperature for 5 minutes, followed by quenching with 250mM glycine at room temperature for 5 minutes. Fixed nuclei were then spun down and washed once with 50% lysis buffer and 50% sucrose buffer prior to ultracentrifugation. Nuclei were resuspended in 1mL 0.1% BSA in PBS and immunostaining was performed.

For bulk RNA-seq, nuclei were unfixed and isolated as described above except that tissue was first dounced in 1mL lysis buffer with 400U RNase inhibitor (Takara, 2313A). Following ultracentrifugation, nuclei were resuspended in 0.1% BSA prior to immunostaining.

For IS-seq, nuclei were isolated as described above. Following ultracentrifugation, they were resuspended in 0.1% BSA in PBS prior to immunostaining.

### Fluorescence in situ hybridization

For DNA FISH of postmortem brain, nuclei extraction was performed as described above. The nuclei pellet was resuspended and fixed in 4% formaldehyde for 10 minutes at room temperature, followed by purification in DPBS. For staining of J-Lat cells, the cells were fixed with 4% formaldehyde for 10 minutes at room temperature and resuspended in DPBS. Approximately 100 µL of either nuclei or cell solution was pipetted onto a charged glass slide before incubation at 60°C for 10 minutes to dehydrate the nuclei. After a brief wash with DPBS, the slides were processed per the RNAscope Multiplex Fluorescent v2 protocol (Advanced Cell Diagnostics). Nuclei were treated with protease, hydrogen peroxide, and incubated with sense DNA probes against the HIV gag-pol gene sequence for 2 hours at 40°C. Amplifier sequences were polymerized to the probes and treated with Opal 570 dye. Nuclei were briefly incubated in DAPI and mounted in VECTASHIELD antifade medium. Imaging was performed at 63x magnification on a Zeiss LSM780 confocal microscope.

For RNA FISH, human cingulate cortex (approximately 100 mg block) from HIV+ and HIV-tissue was drop-fixed in 4% formaldehyde for 6 hours at 4°C, followed by overnight incubation in 30% sucrose. Sections (20 µm) were cut on a freezing microtome and mounted on charged slides. The tissue was processed as described for DNA FISH, with the exception of a longer protease incubation. Tissue was incubated with an Iba1 antibody (ab178846, Abcam) overnight at 4°C and the following day an Alexa-647 goat anti-rabbit (A-21245, Invitrogen) antibody was incubated for 1 hour at room temperature. Prior to mounting in VECTASHIELD, the slides were incubated for 30 seconds in TrueBlack Lipofuscin Autofluorescence Quencher (Biotium) and rinsed with 1x PBS. Images were acquired at 20x and 63x on a Zeiss LSM780 Confocal microscope.

### snRNA-seq

#### Library preparation

snRNA-seq was performed using the Chromium platform (10x Genomics, Pleasanton, CA) with the 3’ gene expression (3’ GEX) V3.1 kit, using an input of ∼16,500 cells. Libraries were sequenced in paired end mode on the NovaSeq (Illumina, San Diego, CA) at an average depth of 176,000 reads per nucleus (range 24,000-200,000).

#### Data processing

Sequencing data was aligned to a combined pre-mRNA GRCh38 and HXB2 reference genome using the Cell Ranger Single-Cell Software Suite (version 3.0.0, 10x Genomics). Filtered count matrices were loaded into R and empty droplets were detected and removed from each sample with Debris Identification using Expectation Maximization (DIEM) (*66*). Doublets were detected and removed using DoubletFinder (*67*). After inspection, two samples which were of low quality were removed (mhbb 154, mhbb 78). This resulted in 69,843 nuclei from 13 samples that passed quality control.

Seurat was used to perform normalization and variance stabilization for each sample prior to merging (*68*). We used Harmony to correct for donor effects (*69*). UMAP dimensional reduction and graph-based clustering were performed on the harmonized dataset in Seurat. After inspection, one cluster that contained low quality nuclei (low read counts and high percent of mitochondrial reads) was removed. Cell cluster identity was assigned based on canonical marker gene expression. We performed further sub-clustering of the microglia cluster. After sub-setting to the cells of interest, HIV was removed as a gene from the dataset. We applied Seurat and Harmony as described above for the full dataset to correct for donor effects and perform graph-based clustering. A cluster that appeared to contain doublets was identified and removed.

For visualization of gene expression, read counts were normalized for each nucleus using log normalization (ln(1+feature count/total counts)). Log normalized counts were then used to generate plots using Seurat.

#### Differential expression analysis

Genes lacking expression in fewer than 10 cells were removed and HIV was removed as a gene from the data. Read counts were summed within each donor for each cluster to create pseudobulk counts. To determine cluster marker gene expression, DESeq2 was applied with a design of ∼ 0 + cluster (*70*). Each cluster was then compared to all other clusters to determine marker genes. For differential expression between conditions (HIV-, HIV+, HIVE), the count matrix was split into sub-matrices for each cluster. We then used DESeq2 with a model of ∼ 0 + condition and identified DEGs by contrasting one condition from another. Differential expression analysis of HIV RNA+ and RNA-microglia was performed similarly. Pseudobulk count matrices were generated within each donor for RNA+ and RNA-microglia and DESeq2 was used with a model of ∼ HIV RNA. Gene ontology enrichment was performed using ShinyGO.(*71*)

#### Analysis of mixed human and mouse snRNA-seq

Samples were aligned to a combined mm10, GRCh38, and HXB2 reference genome. For each nucleus, the proportions of mm10- and GRCh38-mapping reads were calculated and nuclei with >60% of reads mapping to a single genome were considered to be arising from that species. Human nuclei were then filtered using DIEM and DoubletFinder prior to being integrated with the other samples.

For determining HIV read thresholding, nuclei with <600 or >3000 genes detected and >3% of reads mapping to human or mouse mitochondrial genes were removed. Nuclei were identified as human or mouse as described above, and the number of reads mapping to HXB2 in mouse nuclei was determined. For visualization, samples underwent normalization and variance stabilization using SCTransform (Seurat) followed dimensional reduction using UMAP as described above.

#### Immunostaining and fluorescence-activated nuclei sorting

For Hi-C, fixed nuclei were stained with NeuN-488 (MAB377X, Millipore; 1:1000) and Irf8-PE (U31-644, BD Biosciences, 1:250) antibodies. Nuclei sorted for bulk RNA-seq were stained with NeuN-488, Irf8-PE, and Olig2-647 (ab225100, Abcam, 1:500). For IS-seq, nuclei were stained with NeuN-488 (MAB377X, Millipore; 1:1000). All antibodies were incubated for 1 hour at 4°C protected from light. DAPI was added at a concentration of 5ug/mL just before FANS. Nuclei were sorted using a BD Biosciences FACSAria II machine. For Hi-C, fixed nuclei were sorted into PBS and were pelleted in PBS with 17% sucrose buffer by spinning at 4,000rpm for 5 minutes at 4°C. Nuclei pellets were flash frozen on dry ice and stored at -80°C. For bulk RNA-seq, nuclei were sorted directly into Trizol-LS and snap frozen on dry ice. For IS-seq, nuclei were pelleted in PBS with 17% sucrose buffer prior to lysis and DNA extraction as described below.

#### Bulk RNA-seq library preparation and analysis

RNA was extracted from nuclei following manufacturer instructions for Trizol-LS. The aqueous phase containing RNA was then further purified using the RNA Clean & Concentrator kit (Zymo, R1013). RNA-seq libraries were prepared using the SMARTer Stranded Total RNA-Seq Kit v2 Pico Input Mammalian (Takara, 634412). Libraries were sequenced on the Illumina MiSeq platform. Shallow sequencing was used as the aim was to confirm cell-type identity. Following sequencing, reads were trimmed using the default settings of TrimGalore (https://github.com/FelixKrueger/TrimGalore) and aligned to GRCh38 using STAR (settings -- sjdbOverhang 100 --twopassMode Basic) (*72*). Reads were counted using featureCounts (settings --ignoreDup -t ’gene’ -p -O) (*73*). Read counts were log normalized using the DESeq2 rlog function and log normalized read counts were used to generate heatmaps of marker gene expression (*70*).

### Hi-C

#### Library preparation

Nuclei pellets frozen post-FACS were thawed and Hi-C was performed using the Arima-HiC Kit (A51008, San Diego, CA) according to the manufacturer’s instructions. Following enrichment of biotinylated fragments and DNA purification, end repair and adaptor ligation was performed using the Swift Biosciences® Accel-NGS® 2S Plus DNA Library Kit (San Diego, CA), according to the manufacturer’s instructions. DNA libraries were amplified with the Kapa Hyper Prep Kit (NC0709851, Wilmington, MA). Libraries were sequenced on the Illumina NovaSeq platform to an average depth of ∼212 million valid cis-chimeric reads per sample (San Diego, CA, ***Supplementary Table 2b***)

#### Sequencing alignment and contact map generation

We applied HiC-Pro (v2.11.4) to generate contact matrices of HIV- and HIVE microglia (*74*). Briefly, Bowtie2 (parameters: *--very-sensitive -L 30 --score-min L,-0.6,-0.2 --end-to-end -- reorder*) was used to align Hi-C reads to hg38. Uniquely mapped read pairs were assigned to restriction enzyme fragments (DpnII, Hinf1; *ligation sites = GATCGATC, GANTGATC, GANTANTC, GATCANTC*) to filter out invalid ligation products. The resulting valid pairs were then used to generate Hi-C contact maps at 10kb, 40kb, and 100kb resolutions. Hi-C contact maps were normalized using Iterative Correction and Eigenvector decomposition (ICE) via HiC-Pro. Pooled contact maps for HIV-microglia and HIVE microglia were used for the downstream analysis to maximize read depths. We used plotgardener to visualize Hi-C datasets (e.g. contact maps, insulation scores, loops) (*75*).

#### Compartment analysis

We applied dcHiC (https://github.com/ay-lab/dcHiC.git) to HIVE and HIV-microglial contact maps at a 100kb resolution. The resulting output contains genomic coordinates, principal component (PC) values for HIVE and HIV-microglia, and significance (p-values and adjusted p-values) of compartment changes between HIVE and HIV-microglia. Genomic regions with positive and negative PC values were classified as compartment A and B, respectively. We also identified regions with differential compartment signatures between HIVE and HIV-microglia using a threshold of padj< 0.05. Differential compartments were further classified into (1) those that switched compartments (e.g. A in HIVE and B in HIV-) and (2) those that did not switch compartments (e.g. compartment A in both HIVE and HIV-). For the latter ones, compartments were grouped into (1) highA/B (HIV-) to lowA/B (HIVE) when absolute PC values were smaller in HIVE and (2) lowA/B (HIV-) to highA/B (HIVE) when absolute PC values were larger in HIVE. We intersected differential compartments with gene coordinates (Ensembl v93) to identify genes that are located in regions with differential compartment signatures. We then used ShinyGO to perform gene ontology analysis on genes located in differential compartments.

#### Insulation score analysis

We calculated insulation scores from contact maps at a 10kb resolution using *matrix2insulation.pl* from the Github repository: https://github.com/dekkerlab/cworld-dekker.git (parameters: *--is 480000 --ids 320000 --im iqrMean --nt 0 --ss 160000 --yb 1.5 --nt 0 --bmoe 0*). Insulation scores were then converted to TAD boundaries by *insulation2tads.pl* in the same Github repository. For each 10kb bin, the absolute difference in insulation was calculated and the bins with the top and bottom 10% largest changes in insulation score were identified. Data were aligned to hg38 using bwa mem with default settings (*76*). We called peaks using MACS2 with the input ChIP file for background and the settings -B -p .000001 --keep-dup auto (*77*). We performed overlap permutation tests with regioneR to test for overlap between ChIP peaks and insulation score change regions (n=1000 permutation) (*78*).

#### Loop analysis

Loops were called using mustache from contact maps at 10kb resolution of three conditions: HIVE, HIV-, and condition-agnostic (we merged contact maps of HIVE and HIV- to create a chromatin contact map with a higher read depth and resolution) (71). HIVE and HIV- specific loops were identified using the setdiff function in R. To identify genes in HIVE and HIV- specific loops, the entire span of the genome contained within the loop was used. To perform differential loop analysis, we combined loops from HIVE, HIV-, and condition-agnostic. We queried the raw contact frequency of each sample at loop anchors, normalized contact frequencies with the read depth (sum of counts at loop anchors for each sample), and performed differential loop analysis using a linear regression model. Loops with linear regression p < 0.05 were called as differential loops. Difference in contact frequency at differential loops between HIVE and HIV- was further validated by aggregated peak analysis (APA) using hicpeaks (https://pypi.org/project/hicpeaks/). For analysis of genes in differential loops, we identified genes with either of the 10kb anchor regions.

#### Microglial enhancer identification

We obtained previously published human primary microglia H3K27ac ChIP-seq data from dbGaP (***Supplementary Table 2C***) (*79*). Reads were aligned to hg38 using bwa mem with default settings (*76*). Enhancers were called by MACS2 using a p-value threshold of p<0.0001 and a narrowPeak setting (*77*). Super enhancers were called using Rank Ordering of Super-Enhancers (ROSE) with the parameters -s12500 -t2500 (*80, 81*).

### IS-seq

#### Cell culture

JLat clones 8.4, 9.2, 10.6, and 15.4 were obtained from the NIH HIV Reagent Program (ARP-9847, -9848, -9849, and -9850). They were maintained in RPMI 1640 + 10% FBS + penicillin/streptomycin. Jurkat cells were also maintained in RPMI 1640 + 10% FBS + penicillin/streptomycin. Infection of Jurkat cells was performed with NL-43 GFP (NLGI) donated by Benjamin K. Chen for 48 hours.

#### Library preparation

Pelleted nuclei from FANS were resuspended in nuclear lysis buffer (1X Tris-EDTA pH 8, 0.5% SDS), incubated with 40μg/mL RNaseA (12091-039, Invitrogen) for 15 minutes at 37°C, followed by overnight incubation with 50μg/mL proteinase K at 37°C. Genomic DNA was then isolated using phenol chloroform extraction. DNA from cultured cells was extracted using the Qiagen DNeasy Blood & Tissue Kit (cat. 69504).

Integration site libraries were prepared using a linker-mediated PCR (LM-PCR) approach as previously described (*82*). Briefly, DNA was sonicated to an average of 900bp, end repaired, and the DNA fragments were ligated with linkers. Nested PCR was then performed with one primer complementary to the linker and the other primer complementary to the HIV LTR sequence. Libraries were quantified using the KAPA Library Quantification Kit (KK4873). For each set of libraries prepared, we included a positive control of DNA from infected Jurkat cells and a negative control of DNA from HIV-brain or uninfected Jurkat cells. Positive control libraries were not sequenced, negative control libraries were only sequenced in cases where there was amplification and sufficient library was produced to sequence. Final libraries were submitted for sequencing on either the HiSeq 2500 or NovaSeq platforms (Illumina, San Diego, CA).

#### Bioinformatic identification of IS

UMIs were first extracted from the reads using UMI-tools, and then reads were trimmed for HIV LTR sequence and the linker sequence using Cutadapt. Only read pairs for which both the LTR end and linker end had exact matches for both trimmed sequences continued to downstream analysis. We aligned trimmed reads to hg38 using Bowtie2 (settings -p 4 -I 150 -X 1000) and converted reads into a bed file that contained the coordinates of the ends of read 1 and read 2 in order to encompass the both the site of integration (where the LTR-genome junction is located) and the sonic breakpoint (coordinate where the linker ligated).

For each library, reads for which the LTR end was within 5 bp and the sonic end was within 15 bp were collapsed as PCR duplicates and the total number of duplicate reads was counted. For analysis of JLat clones, sonic ends were collapsed within 5bp of one another due to the high level of clonality. Only integration sites with >1 read supporting them were considered as valid. Clonal integration sites were defined as those for which the integration site was the same but the sonic breakpoint was >15 bp apart or >5bp for JLat clones. Any integration sites found in more than one sample, excluding clonal integration sites that came from 2 different assays performed on the same donor, were removed. Any integration sites mapping to non-standard chromosomes were removed prior to analysis.

#### IS characterization

Brain integration site distributions were compared to 3 different T-cell control sets: 1) a compiled dataset of published *in vivo* T-cell integration sites (76,213,225-228), 2) integration sites we sequenced from infected Jurkat cells, 3) a published *in vivo* T-cell dataset for which sequencing data was available (213) and we analyzed using our pipeline. We annotated genomic features using ChIP Seeker (*83*).

### ChIP-seq

#### Library preparation

Normal healthy control frontal cortex tissue gray matter was used to prepare H3K27me3 (n=3 donors) and H3K9me3 (n=3 donors) ChIP-seq libraries. 200-300mg of tissue was homogenized as described above. For H3K9me3 and CTCF, tissue homogenate was fixed with 1% formaldehyde for 10 minutes and quenched with 2% glycine for another 10 minutes rotating at room temperature. Nuclei were isolated and FANS was performed with the NeuN antibody as described above. For H3K27me and H3K9me3 ChIP, after FANS, one million NeuN-nuclei were pelleted and resuspended in 300µl of micrococcal nuclease digestion buffer (10mM Tris pH 7.5, 4mM MgCl, and 1mM Ca2+), and digested with 2 uL of MNase (0.2U/µl) for 5 minutes at 28°C to obtain mononucleosomes. The reaction was quenched with 50mM EDTA pH 8. Chromatin was released from nuclei with the addition of hypotonic buffer (0.2mM EDTA pH 8, containing PMSF, DTT, and benzamidine.

All ChIP samples were then incubated with the appropriate antibodies (H3K27me3, Cell Signaling C36B11; H3K9me3, Abcam ab8898) at 4°C overnight. The DNA-protein-antibody complexes were captured by Protein A/G Magnetic Beads (Thermo Scientific 88803) by incubation at 4°C for 2h. Magnetic beads were sequentially washed with low-salt buffer, high-salt buffer, and TE buffer. DNA was eluted from the beads, treated with RNase A, and digested with Proteinase K prior to phenol-chloroform extraction and ethanol precipitation. For library preparation, ChIP DNA was end-repaired (End-it DNA Repair kit; Epicentre) and A-tailed (Klenow Exo-minus; Epicentre). Adapters (Illumina) were ligated to the ChIP DNA (Fast-Link kit; Epicentre) and PCR amplified using the Illumina TruSeq ChIP Library Prep Kit. Libraries with the expected size (∼275bp) were submitted to the New York Genomics Center for next-generation Illumina sequencing on the HiSeq 2500.

#### Analysis

A mix of publicly available and newly generated ChIP-seq data were used (see ***Supplementary Table 2C***). Both publicly available and new generated ChIP-seq data were aligned to hg38 using bwa mem and called peaks using MACS2. Libraries were subsampled to a similar sequencing depth. For H3K27ac, H3K4me3 and ATAC narrow peaks were called; for H3K27me3 and H3K9me3 broad peaks were called. In cases where there was >1 sample, consensus peaks across multiple samples were called as those occurring in at least 2 samples. Blacklist regions were removed.

### Statistical analysis

Analyses were performed using R 4.1.2. Statistical tests are indicated in the figure legend. P-values of <0.05 were considered significant for chi-square, ANOVA, and Wilcoxon tests. For snRNA-seq analyses, DESeq2 was used to compare expression across summed, pseudobulk data for comparisons of interest. P-values were obtained using the Wald test and multiple comparison correction was performed using the Benjamini-Hochberg method. For differential expression analysis of HIVE vs. HIV-microglia, genes were considered differentially expressed if they either had an adjusted P-value of <0.05 or if they had a log2(fold change)>1 and an unadjusted p-value<0.05. For differential expression analysis of HIV RNA+ vs HIV RNA-HIVE microglia, genes with an adjusted P-value<0.05 were considered differentially expressed. Gene ontology enrichment utilized the hypergeometric test to derive nominal P-values which were then false discovery rate corrected, and categories with an FDR<0.05 were considered significant. Pearson correlations of differential expression results were performed using the log2(fold change) values from DESeq2 results. Hypergeometric P-values used to compare the overlap between genes sets were obtained using an online tool (http://nemates.org/MA/progs/overlap_stats.html) using a background of 58,219 genes in the human genome. Permutation tests used to study the overlap between genomic coordinates were performed using regioneR with n=1000 permutations and all canonical chromosomes as a genomic background (*78*).

## List of Supplementary Materials

### Supplementary table

Table S1 Patient sample metadata

### Supplementary data files 1-10

Data file S1 Sequencing data summary

Data file S2 snRNA-seq cluster marker genes

Data file S3 Quantification of HIV RNA+ nuclei

Data file S4 Hi-C compartment switching regions

Data file S5 Hi-C compartment switching genes

Data file S6 GO complete results

Data file S7 Differential expression results

Data file S8 Hi-C Loop coordinates

Data file S9 Hi-C Loop genes

Data file S10 HIV integration sites

## Acknowledgements

We thank the participants and staff of the Manhattan HIV Brain Bank for their contributions to this work. We thank members of the Akbarian lab, especially Dr. Lucy Bicks, Dr. Jennifer Blaze, Dr. Liz LaMarca, and Dr. Sergio Espeso-Gil for their insightful suggestions and discussions. We thank Dr. Andrew Chess for providing access to his sequencing equipment. We thank the Icahn School of Medicine at Mount Sinai’s Flow Cytometry Core for providing expertise and guidance on nuclei sorting, the Scientific Computing group at the Icahn School of Medicine at Mount Sinai for computational resources and assistance, and the Genomics Core at the Icahn School of Medicine at Mount Sinai for their assistance with single nucleus RNA-sequencing methods.

## Author contributions

Conceptualization: APJ, SM, SA Methodology: APJ

Software: APJ, AV, BH, HW Formal analysis: APJ, BH, HW

Investigation: APJ, AV, CO, MI, BJ, GBH, TL, JM, BK, SC

Visualization: APJ, BJ, HW

Funding acquisition: BC, SM, SA

Project administration: APJ, SM, SA

Supervision: BC, SM, HW, SA

Writing – original draft: APJ, SA

Writing – review & editing: APJ, SA, SM, HW, BC

## Data and materials availability

All sequencing files will be made available on dbGaP. All code will be made available on GitHub.

---

**Figure S1.**
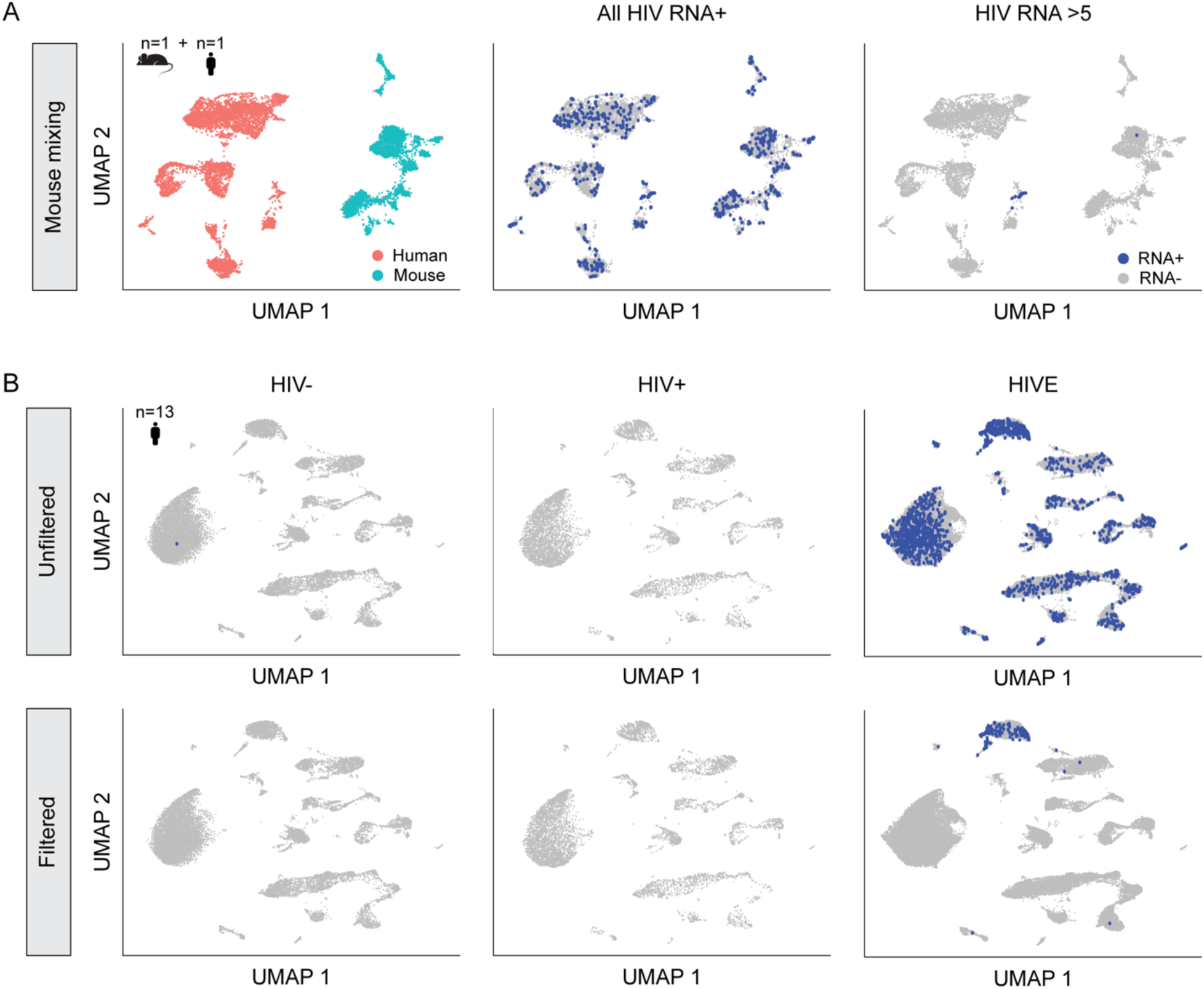
Single nuclei HIV expression thresholding using intermixed mouse-human samples. (**A**) UMAP plot for a representative example of snRNA-seq of mixed mouse and human HIVE tissue showing all HIV RNA+ nuclei and those with >5 HIV reads. (**B**) UMAP plots for the entire snRNA-seq dataset of 13 samples showing HIV RNA expression by sample type for all HIV reads (top row) and only for those nuclei with >5 HIV reads (bottom row).

**Figure S2.**
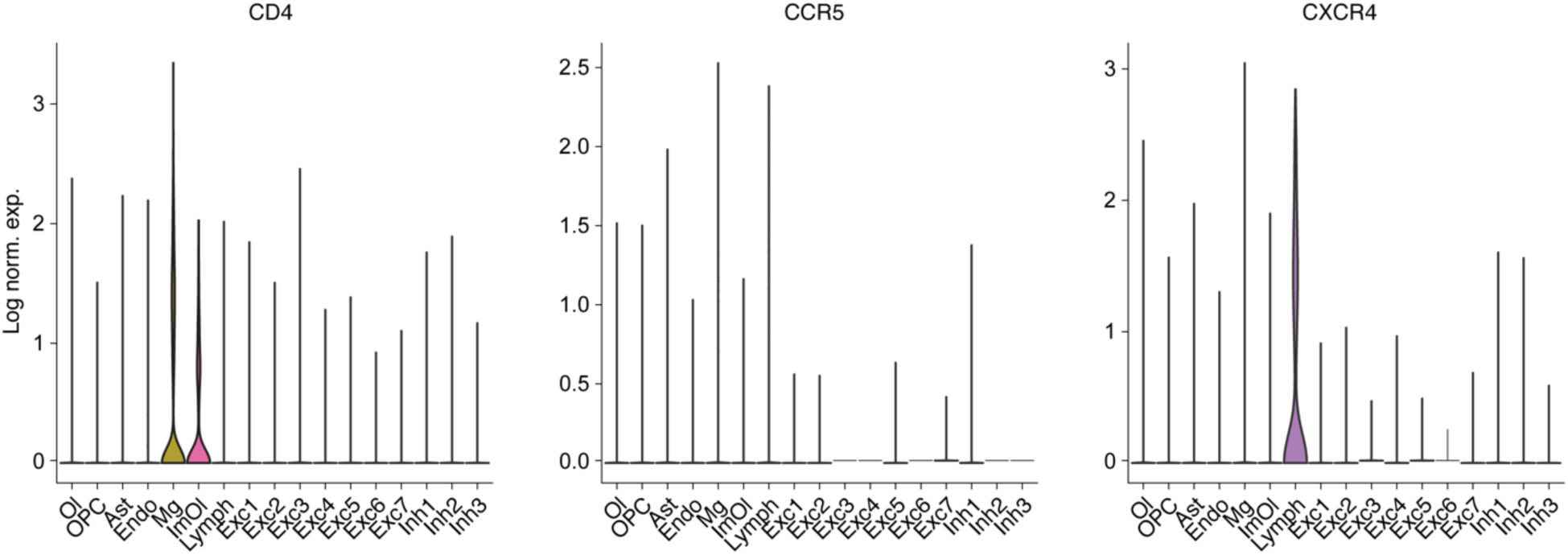
HIV receptor expression in the brain. Violin plots showing expression of CD4, CCR5, and CXCR4 across each cell cluster for the entire snRNA-seq dataset of 13 FC/WM samples.

**Figure S3.**
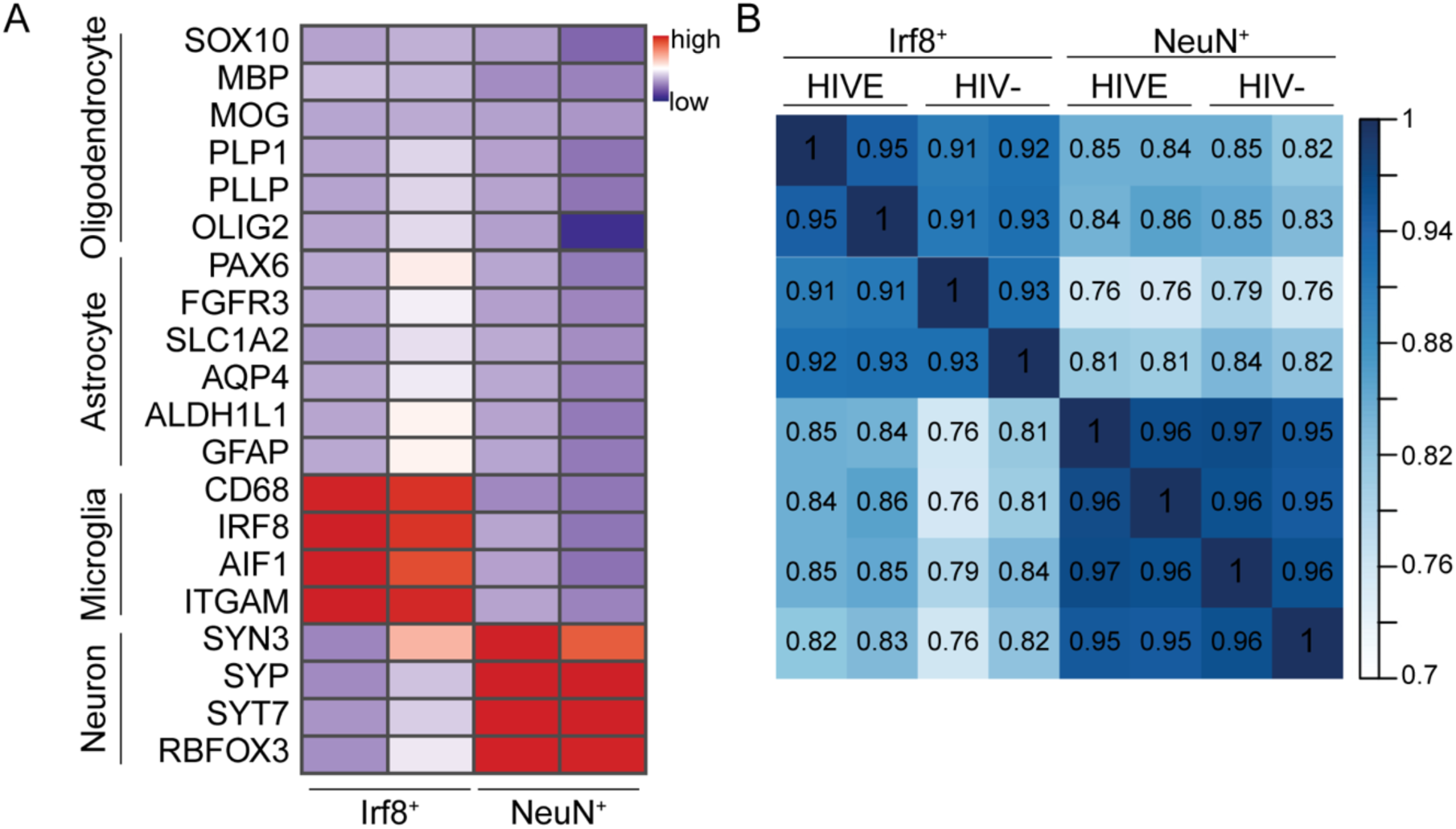
Validation of cell-type-specificity of Hi-C libraries. (**A**) RNA-seq from ensembles of sorted NeuN+ and Irf8+ nuclei showing specific expression of neuronal and microglial marker genes in NeuN+ and Irf8+ nuclei fractions respectively. Heatmap displays scaled transcript per million (TPM) values. (**B**) Sample-by-sample correlations across Hi-C libraries computed using HiCRep.

**Figure S4.**
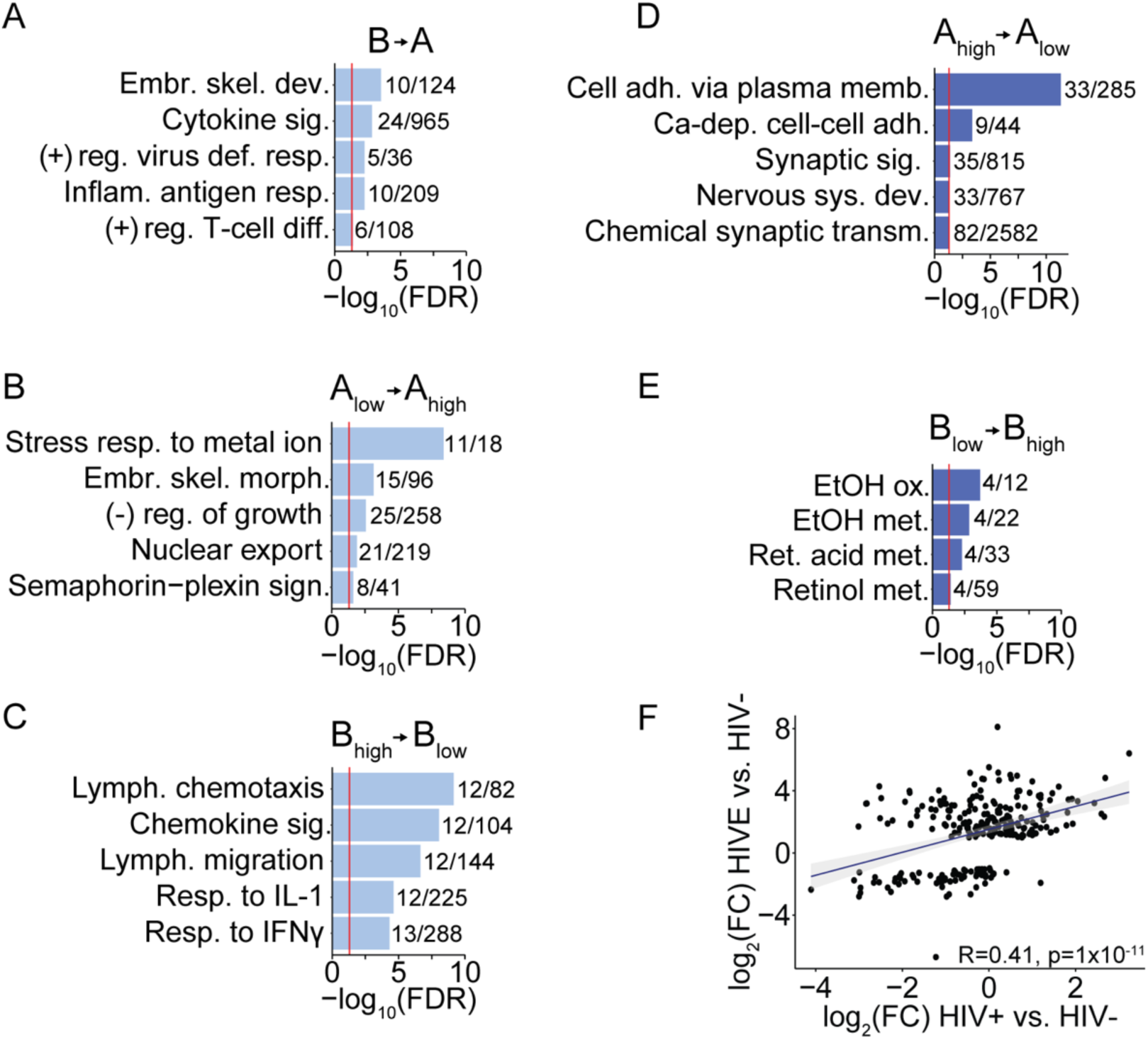
GO enrichment of compartment switching genes. (**A-E**) Gene ontology enrichment for genes in linear genome 10kb bins that had a significant increase (A-C) or decrease (D-E) in PC1 eigenvector value in HIVE microglia compared to HIV-. Specific compartment switches as indicated. Red lines mark significance threshold at FDR=0.05. Numbers indicate the number of genes in the dataset/the number of genes in the corresponding GO category. GO categories: (A) embr. skel. dev., embryonic skeletal development; cytokine sig., cytokine signaling; (+) reg. virus def. resp., positive regulation of virus defense response; inflam. antigen resp., inflammatory antigen response; (+) reg. T-cell diff., positive regulation of T-cell differentiation. (B) Embr. skel. morph., embryonic skeletal morphogenesis; (-) reg. of growth, negative regulation of growth. (C) Lymph. Chemotaxis, lymphocyte chemotaxis; chemokine sig., chemokine signaling; Lymph. migration, lymphocyte migration; resp. to IL-1, response to interleukin 1; resp. to IFNγ, response to interferon-γ. (D) cell adh. via plasma memb., cell adhesion via plasma membrane; Ca-dep. cell-cell adh., calcium-dependent cell-cell adhesion; synaptic sig., synaptic signaling; nervous sys. dev., nervous system development; chemical synaptic transm., chemical synaptic transmission. (E) EtOH ox., ethanol oxidation; EtOH met., ethanol metabolism; ret. Acid met., retinoic acid metabolism; retinol met.; retinol metabolism. Genes undergoing an A→B switch had no significant GO enrichment. (**F**) Pearson correlation between the log2(fold change) in expression of DEGs (shown in Figure 2I) of HIVE vs. HIV-microglia (Y-axis) and HIV+ vs. HIV-microglia (X-axis).

**Figure S5.**
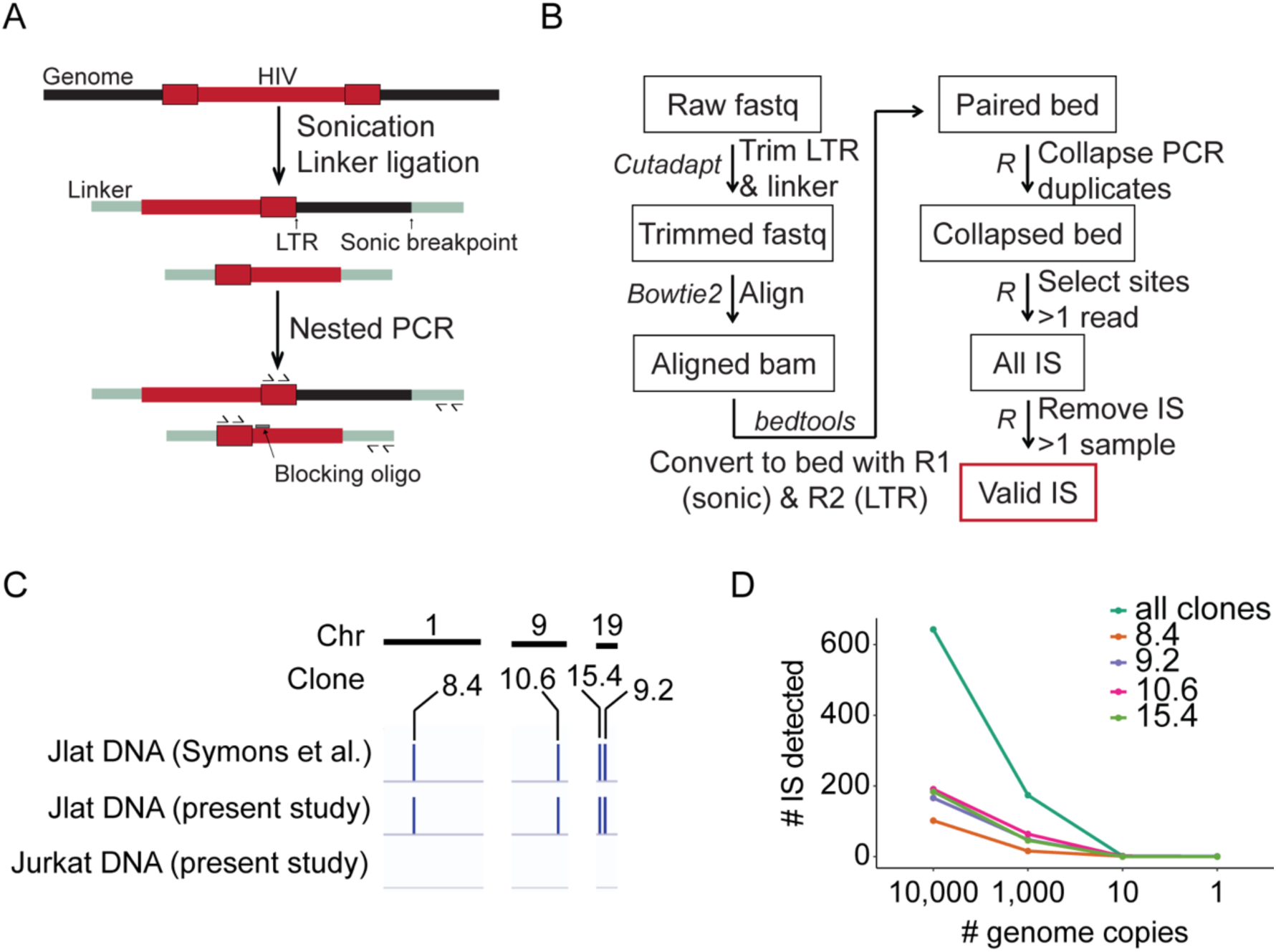
IS-seq specificity, validation, and sensitivity estimation. (**A**) Schematic of integration site library preparation. Sonication and linker ligation was following by nested PCR employing a blocking oligonucleotide to prevent amplification from the 5’LTR. (**B**) Bioinformatic pipeline used for identification of integration sites. (**C**) Browser shots showing results of JLat integration site sequencing compared to previously published JLat sequencing by Symons et al. (*33*). Uninfected Jurkat DNA was used as a negative control. (**D**) Graph showing titration of JLat clone integration site sequencing. The number of JLat genomes input into the library prep is graphed against the number of IS detected.

**Figure S6.**
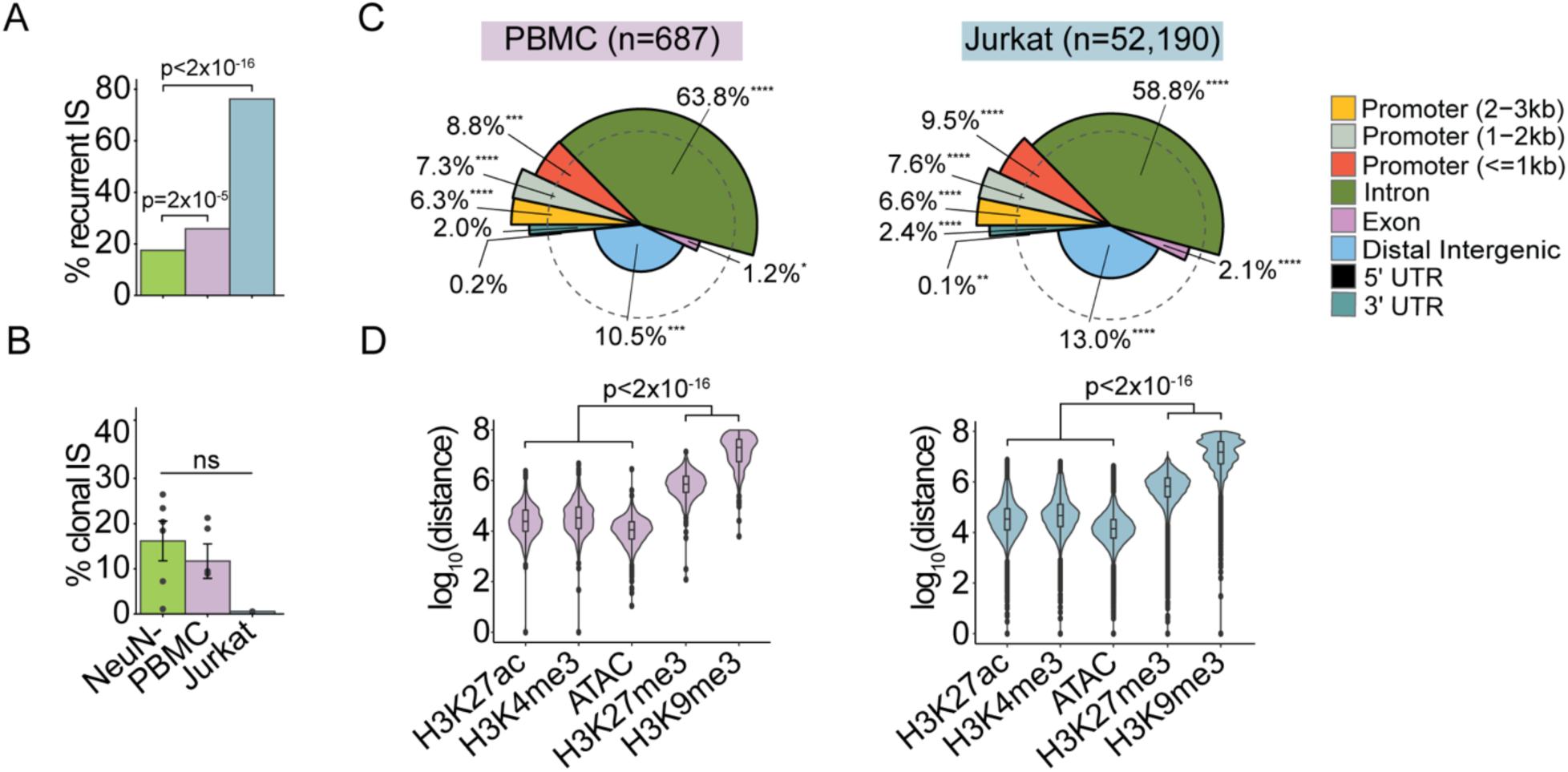
Comparison of NeuN-IS to additional T-cell IS controls. (**A-B**) Percent of IS that are recurrent (A) or clonal (B) for all NeuN-IS, *in vivo* PBMC IS from publicly available IS-seq data (*41*) here analyzed with our pipeline, and integration sites from *in vitro* infected Jurkat cells. Significance by chi-square (A) or two-way ANOVA (B). (**C**) Spie charts comparing the distribution of IS across genomic features to the distribution of those features in the genome. The angle of the pie slice represents the proportion of genome occupied by that feature, while the height of the pie slice represents the relative proportion of that feature in the IS dataset as compared to the genome, dashed circle represents equal proportions. Percentages label the percent of IS found in each genomic feature. Significance using Chi-square test. * 0.01 < p < 0.05, ** 0.001 < p < 0.01, *** 0.0001 < p < 0.001, **** p<0.0001. (**D**) Violin plots showing the distance between IS and the nearest ChIP-seq peak. NeuN- IS are compared to NeuN- ChIP-seq and PBMC and Jurkat datasets are compared to Jurkat ChIP-seq. Significance from Kruskall Wallis rank sum test.

**Figure S7.**
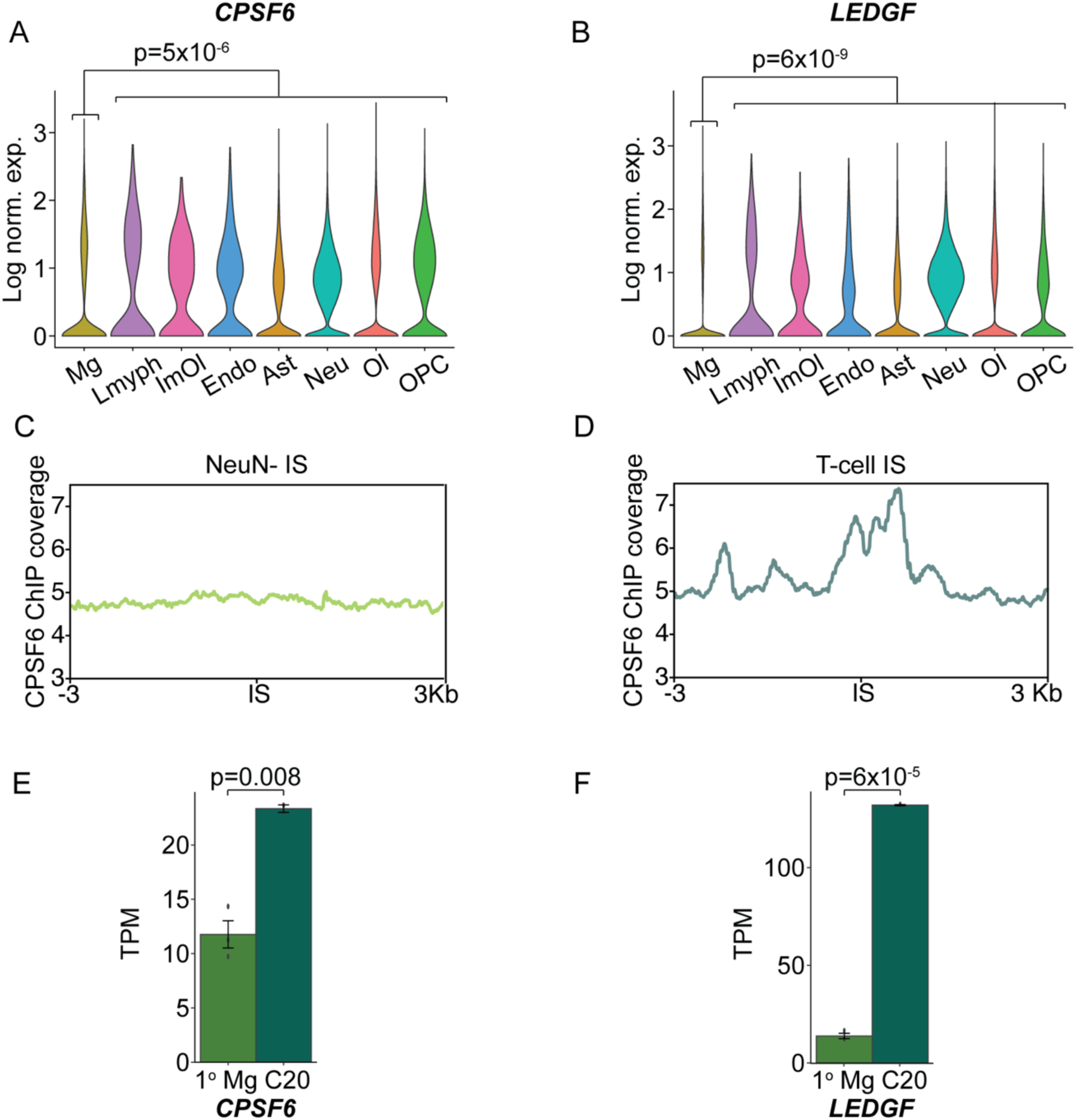
LEDGF and CPSF6 expression in primary microglia. (**A-B**) Violin plots showing expression of *CPSF6* (A) and *LEDGF* (B) from snRNA-seq data for each cell type. For cell types composed of more than 1 subcluster, values were averaged across all cells. P-values reflect DESeq2 differential expression of Mg vs. all other cell types combined. (**C-D**) Bar plots showing transcript per million (TPM) expression of *CPSF6* (C) and *LEDGF* (D) in human adult primary microglia and the C20 human immortalized microglial cell line from a previously published study (PMID **33213476**). Two-tailed Student’s t-test. (**E-F**) Plots showing publicly available CPSF6 ChIP-seq read coverage centered on NeuN- (E) or T-cell (F) IS (PMID 32857953).

**Figure S8.**
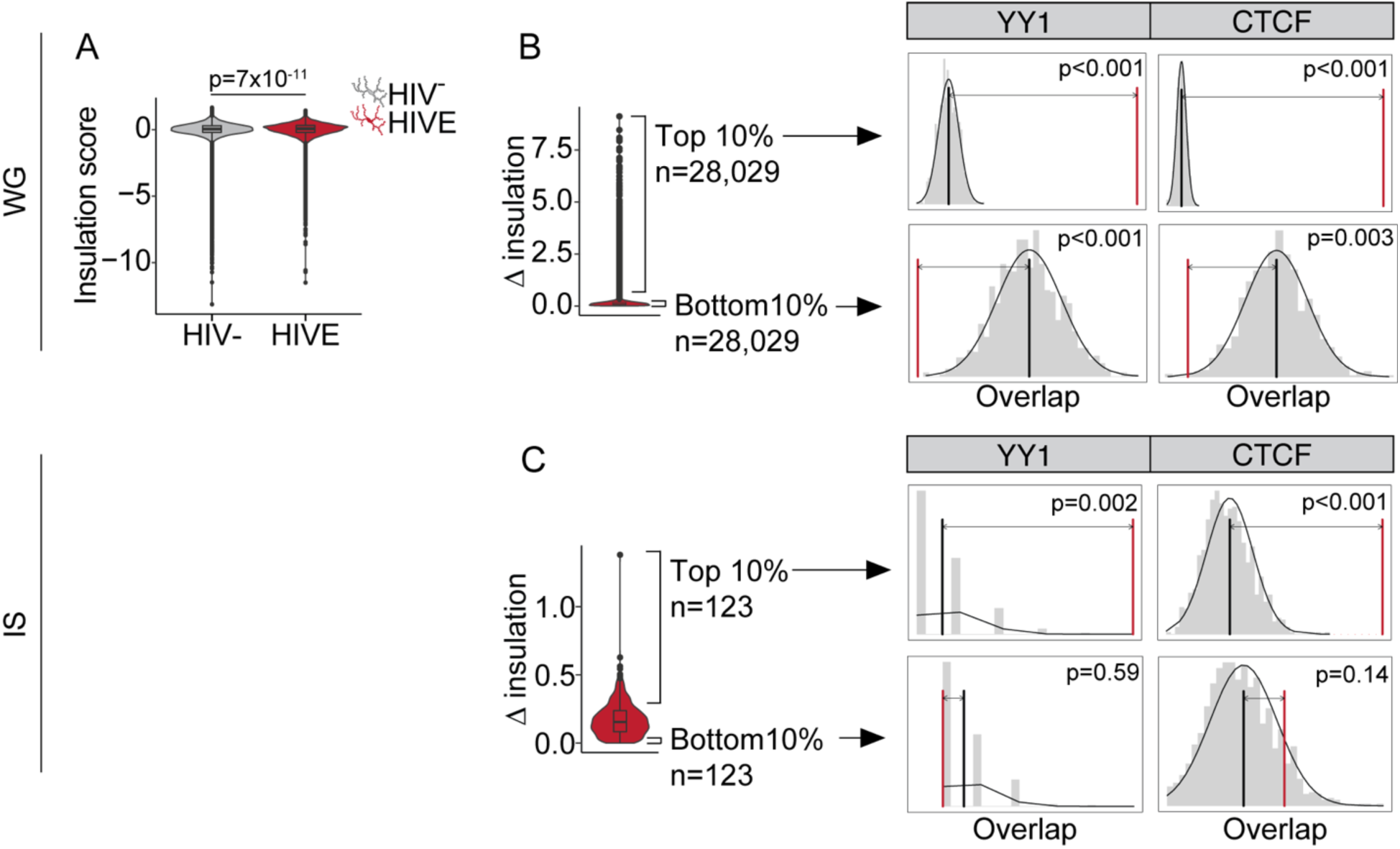
YY1 and CTCF are enriched in regions with large changes in insulation. **(A**) Violin plot showing insulation score across the whole genome (WG) in HIV- and HIVE microglia. Wilcoxon rank sum test. (**B-C**) Change in insulation score between HIVE and HIV- microglia for each 10kb bin of the genome. An overlap permutation (n=1000) test was performed between bins in either top or bottom 10% change in insulation score of the whole genome (B) or just for those bins that contain an IS (C) and ChIP-seq peaks from ENCODE for YY1 and CTCF. Red bar shows the observed number of overlaps between YY1/CTCF ChIP peaks and insulation score regions, black line shows the overlap expected by chance given the whole genome as a background.

**Figure S9.**
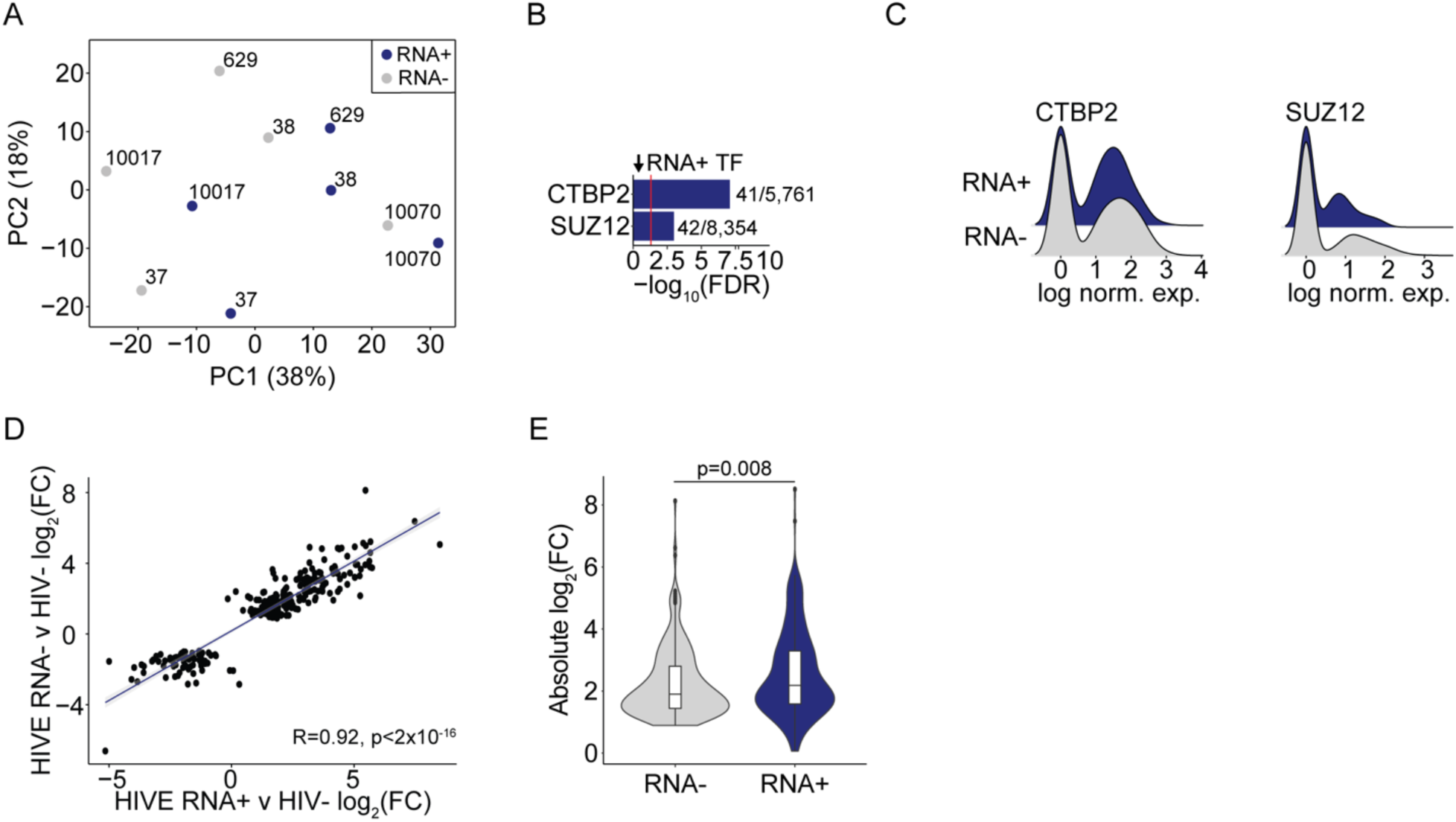
HIV RNA+ microglia are more activated than HIV RNA- microglia. (**A**) PCA of RNA+ and RNA- cells from each HIVE brain with HIV mapping reads present. Donor numbers are indicated. (**B**) Enrichment for ENCODE transcription factor (TF) binding to downregulated genes in HIV RNA+ microglia (shown in Figure 7B). Red line marks significance for FDR=0.05, numbers show the number of differential genes bound by the TF/the total number of genes bound by the TF. (**C**) Ridge plots showing log normalized expression of CTBP2 and SUZ12 in HIV RNA+ and RNA- HIVE microglia. (**D**) Pearson correlation of the log2(Fold Change) in expression between HIV RNA- HIVE Mg vs. HIV- Mg (Y-axis) and the log2(Fold Change) in expression between HIV RNA+ HIVE Mg vs. HIV- Mg (X-axis). Only HIVE vs. HIV- DEGs are shown (from Figure 2I). (**E**) Violin plots showing the absolute log2(Fold Change) for HIV RNA- (gray) or HIV RNA+ (blue) HIVE Mg. Wilcoxon rank sum test.

**Table S1.**
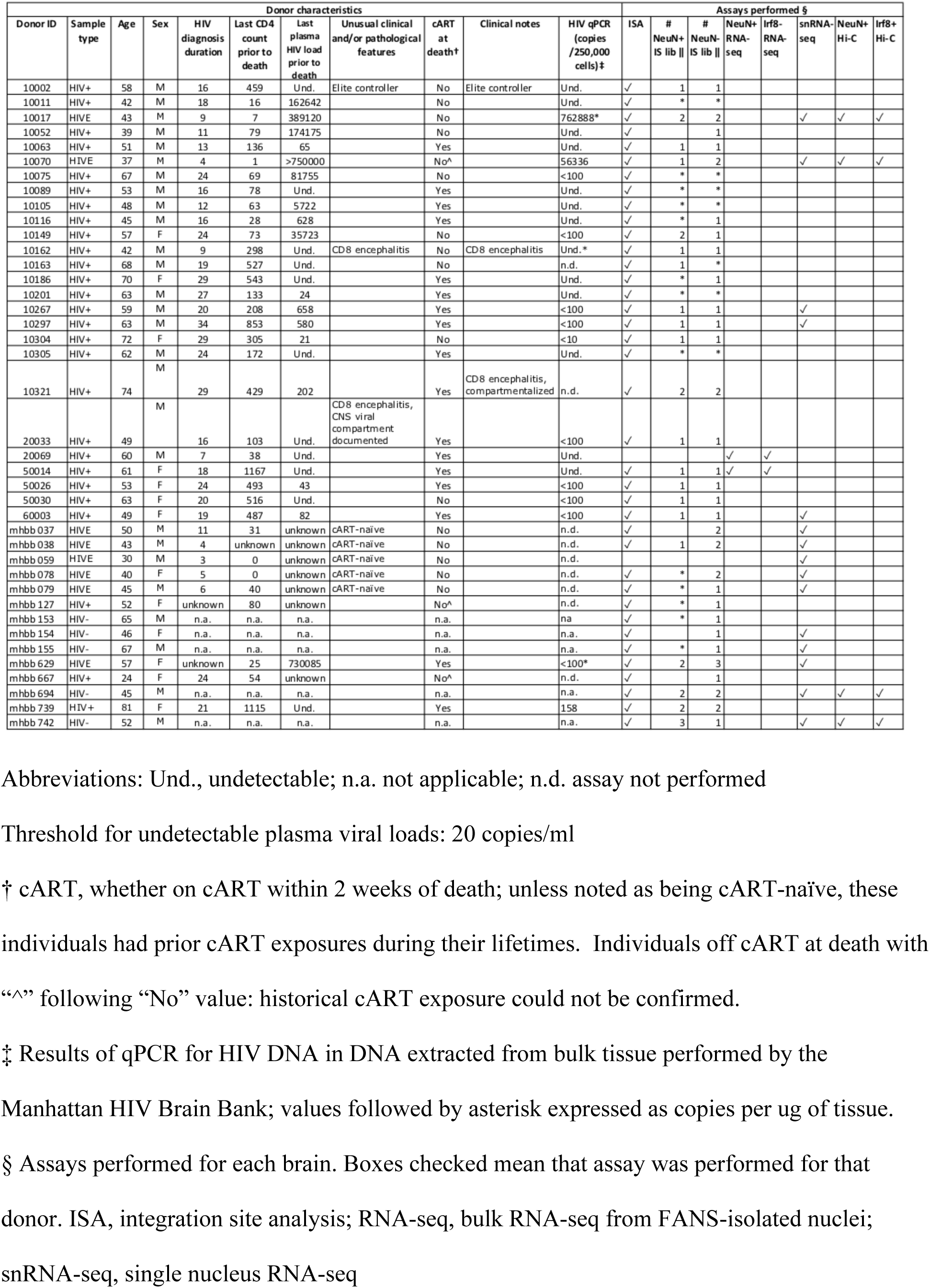
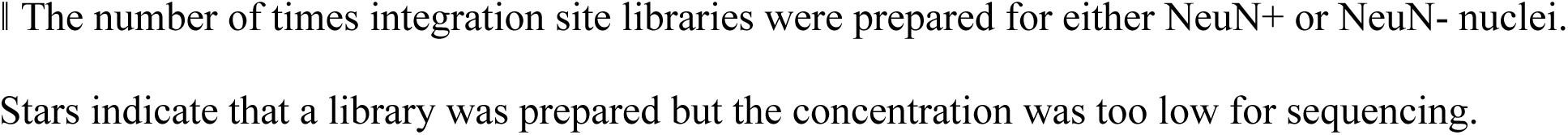
Donor characteristics and assays performed.

